# Verbally specified task goals reorganize visual codes in human ventral temporal cortex via medial frontal modulation

**DOI:** 10.64898/2026.01.24.701461

**Authors:** Robert Kim, Tomas G. Aquino, Sophia Cheng, Chrystal M. Reed, Adam N. Mamelak, Nuttida Rungratsameetaweemana, Ueli Rutishauser

## Abstract

How sensory representations in the sensory cortex are dynamically and rapidly modulated to support flexible, goal-directed behavior remains a fundamental open question. Using rare simultaneous single-neuron recordings across multiple human brain regions, including the ventral temporal cortex (VTC), a high-level visual area not traditionally associated with context-dependent coding, we show that visual representations in VTC are rapidly reconfigured by the presence or absence of preceding verbal task instructions. Identical visual stimuli evoked distinct responses in VTC depending on whether task instructions were provided or not, with categorical representations sharpened when goals were specified in advance. In contrast, dorsal anterior cingulate cortex (dACC) and the hippocampus carried strong signals related to instruction timing, consistent with known regional specialization in top-down control. Coupling between dACC and VTC increased during periods of instructional uncertainty and high cognitive demand on a rapid timescale during stimulus presentation, and its strength predicted correct performance. Together, these findings uncover a rapid, online feedback mechanism through which the medial frontal cortex dynamically reorganizes sensory population codes in VTC, linking flexible cortical coordination to successful goal-directed computation.

## Introduction

Goal-directed behavior requires the brain to dynamically adjust how sensory inputs are represented and interpreted depending on task demands. Human ventral temporal cortex (VTC) has traditionally been viewed as faithfully encoding properties of objects and faces present in the visual input [1–3]. More recently, work in animals and human functional magnetic resonance imaging (fMRI) studies have challenged this view, demonstrating that representations in higher-order visual cortex are not fixed but can be modulated by behavioral state and task context [4, 5]. Despite these advances, the source and circuit mechanisms by which such contextual signals influence visual sensory representations in higher-order visual cortex remain poorly understood, as does the timescale over which they can sculpt ongoing sensory processing. In particular, it is unknown how verbal instructions, the most common way humans acquire goal-related information, shape visual representations in VTC, a region canonically associated with stable visual category coding [1, 6–9], or how rapidly such language-derived goal signals can modulate processing in this area.

Historically, previous studies in both non-human primates and rodents have characterized the ventral visual stream as a primarily feedforward system specialized for encoding stimulus features. In non-human primates, neurons in inferotemporal (IT) cortex, the homologue of the human VTC, exhibit selectivity for visual categories such as faces and objects, forming hierarchical representations that are remarkably invariant to low-level stimulus changes [10–15]. Similarly, neurons in rodent visual areas encode low-level visual features such as orientation, contrast, and shape, with limited sensitivity to behavioral state or task demands [16, 17].

More recent studies utilizing animal models have challenged this purely sensory-encoding view, demonstrating that sensory representations can flexibly change depending on behavioral context, attentional state, or task goals. For example, in rodents, visual cortical responses are modulated by learned associations, expectations, and reward contingencies even when the physical stimuli remain identical [18–23]. In contrast, evidence for such context-dependent modulation in non-human primate visual cortex remains limited. Even less is known in humans, although recent fMRI work has provided evidence that visual cortical representations can be flexibly reorganized by task context [5]. The underlying circuit mechanisms that enable the rapid reorganization of visual representations in response to changing goals in human higher visual areas remain unknown.

We developed a visual decision-making task that is designed to disentangle sensory encoding from modulation driven by the timing of verbal task instructions that either precede or follow the visual stimulus. By combining this task with rare simultaneous single-neuron and local field potential (LFP) recordings from multiple human brain regions spanning the cortical hierarchy, ranging from the VTC to higher-order control areas, we examined how instruction presenece or absence shapes sensory codes at both single-neuron and inter-areal levels. We show that task instructions that precede visual stimuli rapidly modulate sensory codes in human VTC, and further demonstrate that coupling from the dorsal anterior cingulate cortex (dACC) to VTC during sensory processing was critical for successful performance under increased contextual uncertainty and cognitive demand, suggesting that dACC serves as a key source of rapid top-down modulation of VTC. Together, these findings reveal a previously uncharacterized mechanism at the human single-neuron and LFP level through which higher-order control regions dynamically recalibrate sensory representations on behaviorally relevant timescales to optimize goal-directed perception and behavior.

## Results

### Single-neuron recordings across multiple brain areas during a flexible visual decision--making task

We recorded activity from single neurons (773 neurons) in six brain areas, including ventral temporal cortex (VTC; *n* = 164), dorsal anterior cingulate cortex (dACC; *n* = 74), ventromedial prefrontal cortex (vmPFC; *n* = 70), pre-supplementary motor area (preSMA; *n* = 185), hippocampus (HIPP; *n* = 137), and amygdala (AMY; *n* = 143), from 11 patients with refractory epilepsy who underwent depth electrode implantation for clinical monitoring (Fig.1b; see Extended Data Table 1).

**Fig. 1.**
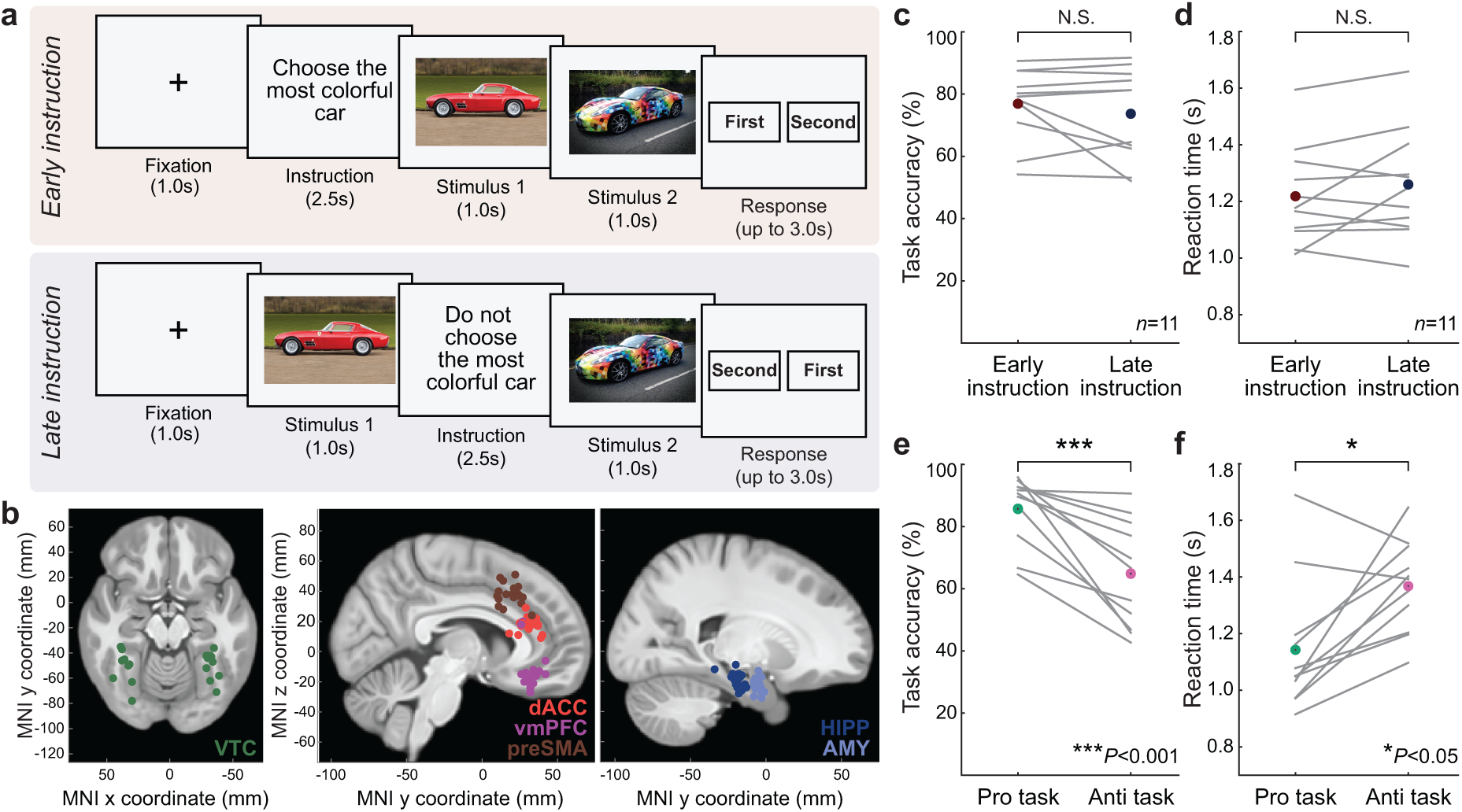
Task design, electrode locations, and behavior. **a**, Two example trials from the flexible visual decision-making task. On each trial, two stimuli were presented sequentially. An instruction was given either before the first stimulus (early instruction; top) or after the first stimulus (late instruction; bottom), specifying whether participants should follow the rule directly (pro) or apply the opposite rule (anti) when making their choice. **b**, Electrode coverage. Colored dots represent microwire bundle electrode locations in Montreal Neurological Institute (MNI) space. **c–f**, Behavioral performance. Task accuracy (**c** and **e**) and reaction times (**d** and **f**) are shown for early vs. late instruction timing conditions (**c** and **d**) and for pro vs. anti tasks (**e** and **f**). Participants showed significantly higher accuracy and faster reaction times in the pro compared to the anti condition, while no significant differences were observed between early and late instruction timing conditions. \**P <* 0.01 and \*\*\**P <* 0.001 by two-sided Wilcoxon rank-sum test; N.S., not significant. VTC, ventral temporal cortex; dACC, dorsal anterior cingulate cortex; vmPFC, ventromedial prefrontal cortex; preSMA, pre-supplementary motor area; HIPP, hippocampus; AMY, amygdala.

Participants performed a flexible visual decision-making task designed to dissociate sensory input from task context (Fig. 1a). On each trial, two visual stimuli were presented sequentially, and participants were instructed either before (“early” instruction) or after (“late” instruction) the first stimulus how to choose between the two (a manipulation we refer to as “instruction timing”). Instructions specified either a “pro” condition, in which participants followed the specified rule directly, or an “anti” condition, in which they applied the opposite rule (see Extended Data Table 2 for examples instructions). The anti condition imposed greater cognitive demands due to its double-negative structure. Crucially, the same images were presented in both early and late instruction conditions, enabling us to investigate how VTC and other regions represent identical visual inputs under different task contexts.

Behaviorally, participants performed the task with high accuracy (mean *±* s.d.; 75.28 *±* 12.45%, chance is 50%). Accuracy was significantly higher for pro compared to anti trials, consistent with greater cognitive demands in the anti condition (Fig.1c; mean *±* s.d., 85.70 *±* 11.14% vs. 64.87 *±* 17.04%; *P <* 0.001 by two-sided Wilcoxon signed-rank test). Reaction times were also longer for anti compared to pro trials (mean *±* s.d.; 1.41 *±* 0.16s vs. 1.17 *±* 0.23s; *P <* 0.005 by two-sided Wilcoxon signed-rank test). In contrast, no significant differences in accuracy or reaction times were observed between early and late instruction trials (accuracy: 76.14 *±* 12.34% for early vs. 74.43 *±* 14.30% for late; reaction time: 1.29 *±* 0.16s vs. 1.29 *±* 0.20s; *P >* 0.5, two-sided Wilcoxon signed-rank test). To further examine behavioral effects, we compared early versus late instruction trials within pro and anti conditions separately. Although task performance did not differ significantly between early and late trials within either the pro or anti conditions, early pro trials showed significantly faster reaction times than late pro trials (*P <* 0.05 by one-sided Wilcoxon signed-rank test; Extended Data Fig. 1). No significant difference in reaction time was observed between early and late trials for the anti condition (Fig. 1d).

### VTC neurons are modulated by both stimulus category and instruction timing

We first asked whether single-neuron activity in human VTC reflects not only the visual category of the stimulus but also the timing of task instruction, which determines when the goal is known relative to stimulus onset. We first examined the firing rates of single-neurons following onset of stimulus 1 (stim 1). We observed robust category selectivity in many VTC neurons (examples shown in Fig. 2a,b), consistent with the established role of VTC in forming sensory representations of visual categories [24].

**Fig. 2.**
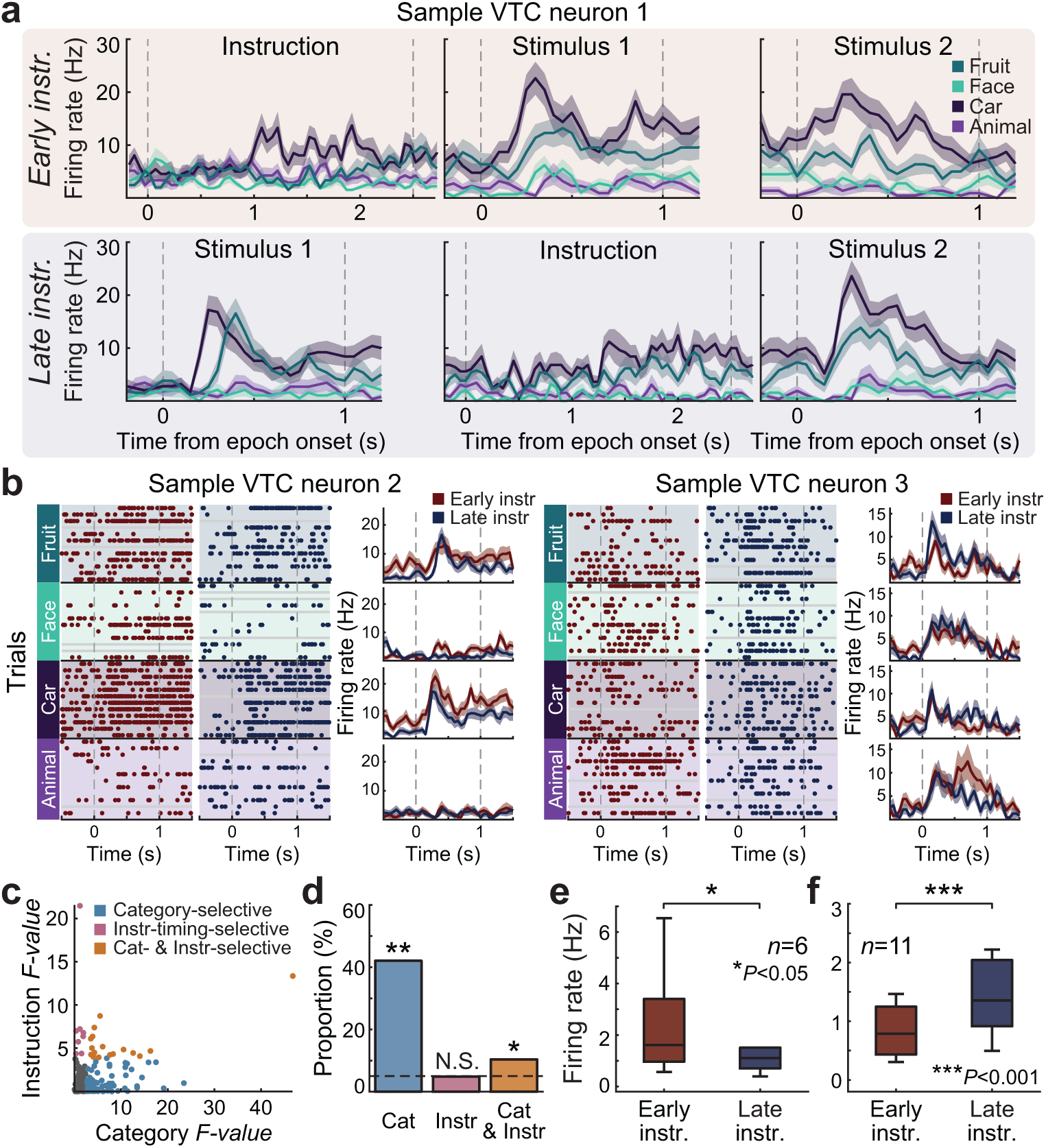
VTC neurons encode stimulus category and instruction timing. **a**, Example VTC neuron showing category-selective responses (to car images in this case) during both the stimulus and instruction windows. **b**, Two additional sample VTC neurons illustrating instruction-related modulation. Neuron 2 showed enhanced responses to preferred categories during early instruction trials, while Neuron 3 showed diminished responses under the same condition. Raster plots (left) and peristimulus time histograms (right) are aligned to stimulus 1 onset. Each row in the raster plot corresponds to a single image, shown under both early and late instruction conditions. **c**, Two-way ANOVA (category *×* instruction timing) during the stim 1 window revealed three classes of VTC neurons: category-selective (cyan), instruction-timing-selective (magenta), jointly selective (orange), and non-selective (black). **d**, Proportion of neurons in each category, showing that most were category-selective, with a smaller but notable jointly selective population. The dashed line indicates the 5% chance level. \**P <* 0.005 and \*\**P <* 0.001 by binomial test; N.S., not significant. **e, f**, Average firing rates during the stim 1 window for jointly selective neurons. These neurons could be divided into two groups: one subgroup showed enhanced responses to their preferred category during early instruction trials (**e**), while the other subgroup showed suppressed responses under the same condition (**f**). Boxplot central lines, median; bottom and top edges, lower and upper quartiles; whiskers, 1.5*interquartile range; outliers not plotted. \**P <* 0.01 and \*\*\**P <* 0.001 by two-sided Wilcoxon signed-rank test.

Performing a 4 *×* 2 two-way ANOVA with interactions with factors of visual category (there were four possible categories) and instruction timing during the stim 1 window revealed a large proportion of VTC neurons that were category-selective (69 of 164 neurons, 42.07%, *P <* 0.001 by binomial test; Fig. 2c,d). The preferred category was determined as the category that elicited the highest mean firing rate during the stim 1 window for the early instruction trials only. In contrast, only 8 neurons (4.9%, *P >* 0.5 by binomial test) exhibited a significant main effect of instruction timing in the absence of a significant category effect, indicating that contextual modulation alone was uncommon. A larger subgroup of neurons exhibited significant main effects of both stimulus category and instruction timing (e.g., joint selectivity), with category responses that varied depending on whether the instruction was presented early or late (Fig. 2c,d; 17 of 164 neurons, 10.37%, *P <* 0.005 by binomial test). This proportion was also greater than expected if category and instruction selectivity were independent (*χ*^2^(1) = 8.14, *P* = 0.0037). Within this jointly selective subpopulation, some neurons showed enhanced responses to their preferred category during early instruction trials, whereas others showed suppressed responses under the same condition (Fig. 2e). These results demonstrate that while category selectivity dominated in VTC, a subset of neurons flexibly integrated both sensory and contextual information, suggesting that VTC may be subject to context-dependent top-down modulation.

Several of the category-selective neurons were also active during the instruction period when their preferred category was mentioned verbally, for early (7 of 57 category-selective VTC neurons) and late instruction (15 of 57) trials, suggesting that verbal task cues can activate or sustain category-specific representations in VTC (see Extended Data Fig. 2). In addition, some neurons exhibited pronounced instruction-dependent modulation of category responses: activity was either enhanced (Fig. 2b, left) or suppressed (Fig. 2b, right) when the instruction preceded stimulus 1, indicating flexible gain control dependent on task context.

### Population-level decoding of instruction timing and category in VTC

Given the modulation related to instruction timing we observed at the single-neuron level in VTC, we next asked whether we could reliably decode the presence of instruction during the stimulus windows from VTC population activity. Using a linear SVM decoder during the stim 1 window, we found that VTC population activity (pseudopopulation of all recorded neurons across participants) reliably differentiated early from late instruction trials, with decoding accuracy significantly above chance (Fig. 3a; mean *±* s.d., 64.78 *±* 6.68% vs. 50.66 *±* 7.12% for shuffled data). We next visualized the structure of VTC population activity by projecting it into a low-dimensional latent space using Contrastive Embedding for Behavioral and Neural Analysis (CEBRA; [25]). CEBRA is a non-linear dimensionality reduction framework designed for uncovering latent population dynamics from neural data. In this analysis, we used an unsupervised implementation (i.e., without providing any task or behavioral labels) to visualize the intrinsic structure of VTC activity across the two instruction timing conditions (early and late; see Methods). Even without explicit task information, this approach revealed distinct neural trajectories for early and late instruction conditions during the stim 1 window (Fig. 3b). Computing the distance between the two trajectories revealed that early and late instruction trials were reliably separated during the stim 1 period compared to shuffled controls (Fig. 3c).

**Fig. 3.**
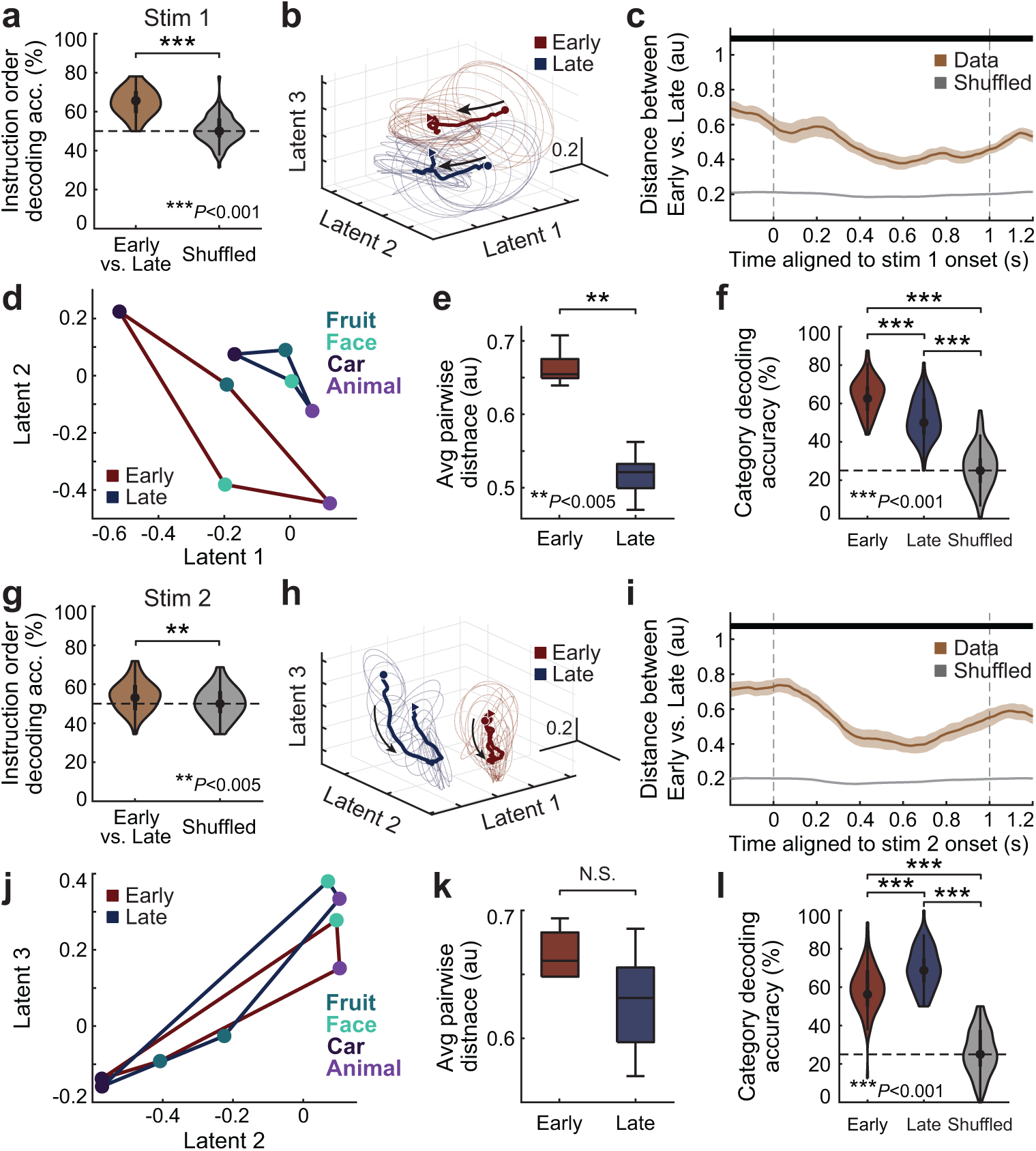
VTC population activity encodes instruction timing and sharpens category representations. **a**, Linear SVM decoding during the stim 1 window using a pseudopopulation of all recorded VTC neurons revealed that instruction timing (early vs. late) could be decoded significantly above chance. **b**, Low-dimensional neural trajectories obtained with CEBRA for early (blue) and late (red) instruction trials during the stim 1 window. **c**, Distance between early and late trajectories shown in b, compared with shuffled controls. **d**, Category trajectories in the CEBRA latent space for early versus late instruction trials during the stim 1 window (at 0.70 s after stim 1 onset). **e**, Average pairwise category distances during the stim 1 window. **f**, Linear SVM decoding accuracy for category identity during the stim 1 window for early and late instruction trials. **g-j**, Similar analyses as in **a**–**f**, performed during the stim 2 window. Ellipsoids in the CEBRA trajectories (**b**, **h**) show trial variability, estimated from the covariance of neural activity at selected time points and plotted as orthogonal projections in latent space. Boxplot central lines, median; bottom and top edges, lower and upper quartiles; whiskers, 1.5*interquartile range; outliers not plotted. \*\**P <* 0.005 and \*\*\**P <* 0.001 by two-sided Wilcoxon rank-sum test (**a**, **e**, **g**, **k**) or Kruskal-Wallis test with post hoc Dunn’s multiple comparisons test (**f**, **l**). Thick horizontal black bars in **c** and **i** indicate time periods of significant differences identified using a cluster-based bootstrap procedure (*P <* 0.05, cluster-level corrected; see Methods). Circles and triangles mark the onset and offset of the stim 1 analysis window, respectively.

How did instruction timing information modify representations of categories in VTC? When the instruction was provided before stim 1 (i.e., early instruction trials), category representations were sharpened, as indicated by larger pairwise distances between category trajectories in the CEBRA latent space (Fig. 3d) and greater overall category separation (Fig. 3e). This increase in representational distance suggests that advance knowledge of task goals allows VTC population activity to more effectively differentiate between stimulus categories. In contrast, when the instruction was given after the first stimulus (i.e., late instruction trials), trajectories corresponding to different categories were more overlapping (Fig. 3d, blue), reflecting a less differentiated sensory representation during the initial encoding phase. Consistent with these findings, stimulus category could be decoded with significantly higher accuracy in early compared to late instruction trials (Fig. 3f; mean *±* s.d., 64.62 *±* 8.67% for early, 52.12 *±* 10.52% for late, and 26.75 *±* 10.88% for the shuffled null condition), confirming that contextual information enhances the fidelity of category-specific coding in VTC.

We next performed the same analyses during the stim 2 window. During this time window, instructions have always been shown but in some instances just before stim 2 was shown (late) vs. already before the stim 1 window. By this stage both the early and late conditions thus contained equivalent amounts of sensory and contextual information. Despite this, at the population level, VTC still differed between the early and late conditions (Fig. 3a), albite with less accuracy than during the stim 1 window (Fig. 3a; mean *±* s.d., 53.81 *±* 7.35% vs. 50.50 *±* 7.69% for shuffled data). Neural trajectories in the CEBRA latent space also remained well separated between early and late instruction conditions (Fig. 3h,i), indicating that contextual timing differences in VTC population activity persisted throughout the trial. Pairwise category distances were large for both early and late instruction trials at this stage (Fig. 3j,k), suggesting that category information became robustly represented in VTC regardless of instruction timing. However, decoding again revealed higher category decodability when the instruction immediately preceded the stimulus, corresponding to the stim 2 window for late instruction trials (Fig. 3l; mean *±* s.d., 59.06 *±* 11.80% for early, 69.75 *±* 10.72% for late, and 26.12 *±* 11.53% for the shuffled null condition). Together, these results indicate that VTC population activity flexibly integrates sensory information with instruction timing across time, with category representations sharpened when task context immediately preceded stimulus presentation.

### Signals related to instruction timing in hippocampus and dACC

Having shown that VTC encodes not only stimulus category but also exhibits modulation by instruction timing for the same stimuli, we next investigated potential sources of these top-down signals, defined here as contextual influences from higher-order brain regions that shape sensory responses. Recent animal studies demonstrate that top-down feedback signals from frontal cortex can dynamically modulate visual cortical processing. For example, Ährlund-Richter et al. showed that mouse prefrontal subregions, especially anterior cingulate cortex, exert behavioral-state-dependent feedback onto primary visual cortex [15]. Motivated by these insights, we next asked whether higher-order human brain areas, including hippocampus and dorsal anterior cingulate cortex (dACC), encode instruction timing signals that could provide top-down inputs necessary to flexibly shape VTC sensory representations.

We first quantified single-neuron selectivity in hippocampus and dACC using a 4 *×* 2 two-way ANOVA with factors of category and instruction timing, as in VTC (Fig. 4a). Both regions contained more neurons selective for instruction timing than VTC, with dACC showing the highest proportion (HIPP, 20 of 137 neurons, 14.60%, *P <* 0.001 by binomial test; dACC, 17 of 74 neurons, 22.97%, *P <* 0.001 by binomial test). Decoding similarly followed this hierarchy, with accuracy for distinguishing early versus late instruction trials highest in dACC (Fig. 4b). We also extended the two-way ANOVA analysis to the amygdala (AMY), pre-supplementary motor area (preSMA), and ventromedial prefrontal cortex (vmPFC) (Extended Data Fig. 3). The amygdala showed a pattern similar to the hippocampus, with a moderate proportion of neurons selective for instruction timing. In contrast, both preSMA and vmPFC exhibited patterns resembling dACC, with a larger fraction of neurons showing instruction-related modulation, consistent with recent human single-neuron recordings demonstrating that preSMA neurons encode abstract decision variables and integrated choice signals during flexible behavior [26]. Consistent with this pattern, decoding of instruction timing revealed higher accuracy in medial frontal regions (preSMA, dACC, and vmPFC) compared to VTC during the stim 1 window (Extended Data Fig. 4 left). Although this hierarchical separation was less pronounced during the stim 2 window, medial frontal areas continued to show robust instruction-related signals relative to VTC (Extended Data Fig. 4 right). These findings confirm that instruction timing is represented across a distributed network of higher-order regions, with the strongest contextual modulation emerging in the medial frontal cortical areas dACC, preSMA, and vmPFC.

**Fig. 4.**
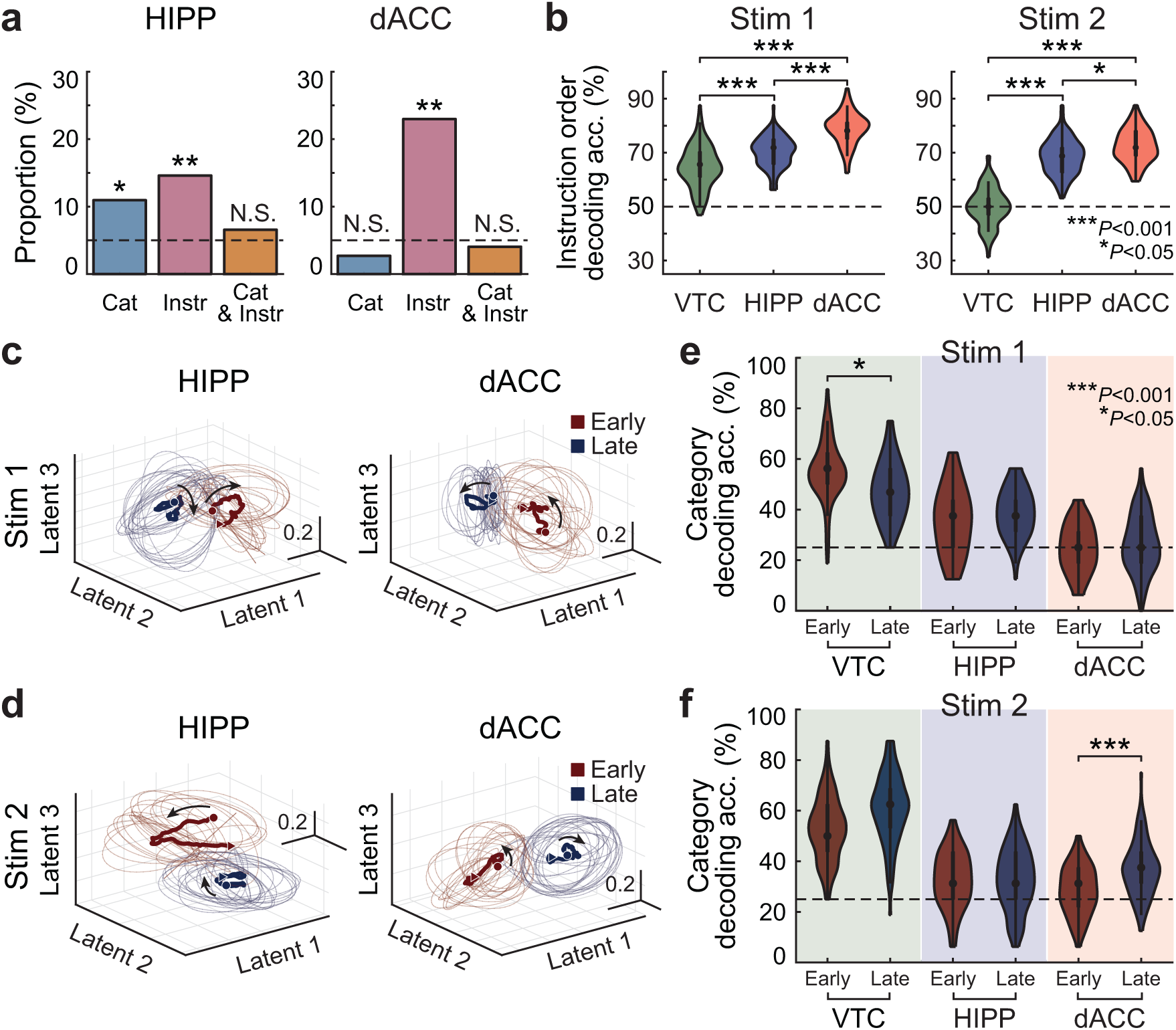
Signals related to instruction timing in hippocampus (HIPP) and dorsal anterior cingulate cortex (dACC). **a**, Proportion of neurons selective for stimulus category (cyan), instruction timing (magenta), or both (orange), based on two-way ANOVA (category *×* instruction timing). \**P <* 0.005 and \*\**P <* 0.001 by binomial test; N.S., not significant. **b**, Linear SVM decoding accuracy for distinguishing early versus late instruction trials in HIPP and dACC during the stim 1 window (left) and stim 2 window (right). **c**, Low-dimensional population trajectories obtained with CEBRA for early (blue) and late (red) instruction trials in HIPP and dACC during the stim 1 window (top) and stim 2 window (bottom). Ellipsoids indicate trial variability estimated from the covariance of trajectories at selected time points. **d**, Linear SVM decoding accuracy for stimulus category across regions, showing highest decoding in VTC and reduced decoding in HIPP and dACC. \**P <* 0.05 and \*\*\**P <* 0.001 by Kruskal-Wallis test with post hoc Dunn’s multiple comparisons test. Circles and triangles mark the onset and offset of the stim 1 analysis window, respectively.

Visualization of population activity using CEBRA revealed that trajectories in both hippocampus and dACC were well separated between early and late instruction conditions, demonstrating robust representation of instruction timing at the population level (Fig. 4c,d).

We then examined how stimulus category information was represented across these regions. Decoding revealed that category decoding ability decreased systematically along the hierarchy, with VTC showing the highest decoding and HIPP and dACC showing lower decoding accuracy; however, decoding in HIPP remained significantly above chance (Fig. 4e,f; Matched for number of neurons across areas). Extended Data Fig. 5 further illustrates this pattern by showing category decoding for early and late instruction trials across all regions. Consistent with the hierarchy observed in the main analysis, VTC exhibited the highest category decodability during both stimulus windows, whereas medial frontal and medial temporal regions showed markedly lower decoding accuracy, reinforcing the distinction between lower-level sensory regions that primarily encode stimulus features and higher-order areas that emphasize task context. VTC again showed sharpened category representations when the instruction immediately preceded the stimulus window of interest. The results here differ slightly from Fig. 3f,l because the number of neurons was matched across regions to the smallest sample size, which was 74 in dACC. This pattern is consistent with VTC’s established role in bottom-up sensory coding, whereas higher-order regions primarily emphasized instruction-related information.

Together, these analyses suggest that higher-order regions such as dACC may provide top-down signals that modulate VTC, enabling sensory representations in VTC to be flexibly shaped by task context.

### dACC-VTC modulation reflects uncertainty, cognitive demand, and behavioral performance

We next employed spike-field coherence (SFC) analyses to examine potential sources of top-down modulation to VTC. Previous studies in non-human primates demonstrated that feedback projections from prefrontal and cingulate areas modulate sensory processing in visual cortices through rhythmic synchronization across broad frequency ranges, including the theta (4-7 Hz) and beta (12-32 Hz) bands [15, 27, 28]. Motivated by these findings, we tested whether VTC spikes phase-locked to ongoing oscillations in the LFP in the other simultaneously recorded areas, including dACC (Fig. 5a; see Methods). Focusing on the stim 1 window and comparing early versus late instruction trials revealed that VTC spikes were more strongly phase-locked to LFPs from dACC across a broad frequency range in early relative to late trials, especially in the beta band (12-32 Hz; Fig. 5b). Investigating whether LFP activity from other regions was coordinated with VTC spiking revealed that the amygdala also exhibited strong late-instruction modulation, primarily within the theta range (Extended Data Fig. 6). Examining the reverse direction, i.e. whether spikes in other areas were phase-locked to VTC LFP, showed robust early-instruction modulation from VTC to dACC, potentially indicating that early contextual information sharpens dACC-VTC interactions (Extended Data Fig. 6).

**Fig. 5.**
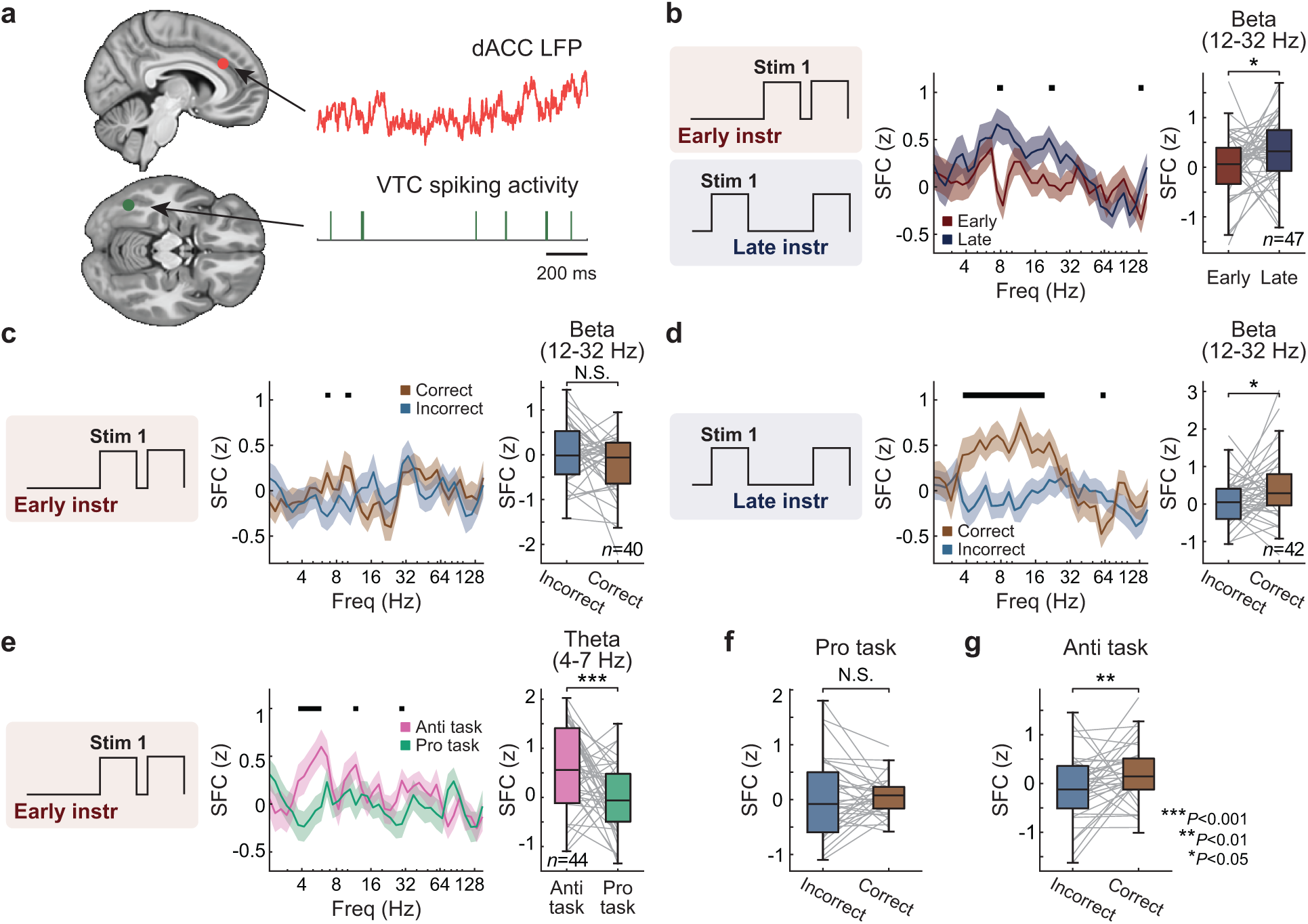
dACC-VTC spike-field coherence (SFC) reflects uncertainty, cognitive demand, and behavioral performance. **a**, Example recording configuration showing simultaneous dACC local field potential (LFP) signal and VTC spiking activity. **b**, SFC aligned to stimulus 1 for early vs. late instruction trials. Left, schematic of trial types. Middle, SFC spectra showing increased beta-band (12–32 Hz) coherence for early compared to late instruction trials. Right, paired comparisons of beta-band SFC across neurons (*n* = 47). **c**, SFC during stimulus 1 for correct vs. incorrect trials. Left, schematic. Middle, spectra showing comparable coherence between correct and incorrect trials. Right, paired comparisons in the beta band (*n* = 40). **d**, SFC during stimulus 1 for correct vs. incorrect trials, showing significantly increased beta-band coherence on correct compared to incorrect trials (*n* = 42). **e**, SFC for pro vs. anti trials. Left, schematic. Middle, spectra showing elevated theta-band (4–7 Hz) coherence in anti compared to pro trials. Right, paired comparisons of average theta-band SFC (*n* = 44). **f**, Average theta-band SFC for correct vs. incorrect trials separated by task condition, shown for pro (left) and anti (right) trials. Thick horizontal black bars indicate time periods of significant differences identified using a cluster-based bootstrap procedure (*P <* 0.05, cluster-level corrected). Boxplot central lines, median; bottom and top edges, lower and upper quartiles; whiskers, 1.5*interquartile range; outliers not plotted. \**P <* 0.05, \*\**P <* 0.01, and \*\*\**P <* 0.001 by two-sided Wilcoxon signed-rank test.

These findings suggest that dACC exerts modulation of VTC during periods of uncertainty, specifically in late instruction trials within the stim 1 window during which instructions have not yet been provided by the time the stimulus appears. We next asked whether this top-down modulation was behaviorally relevant by examining whether the strength of dACC-VTC SFC was associated with task performance. We compared dACC-VTC SFC between correct and incorrect trials, separately for early (Fig. 5c) and late (Fig. 5d) instruction conditions. We ensured that the number of trials and spikes was matched between correct and incorrect sets within each instruction timing condition (see Methods). During early instruction trials, dACC-VTC SFC did not differ significantly between correct and incorrect responses (Fig. 5c). In contrast, during late instruction trials, dACC-VTC coupling in the beta band was significantly stronger on correct compared to incorrect trials (Fig. 5d). Repeating this analysis in the reverse direction (i.e., SFC between VTC LFP and dACC spikes) revealed no significant increases for correct trials (Extended Data Fig. 7), indicating that performance-related modulation was specific to top-down dACC *→* VTC interactions rather than bottom-up signaling. These results indicate that when task uncertainty was high (because instructions had not been provided yet), stronger top-down modulation from dACC to VTC was associated with successful performance. This pattern suggests that, in the absence of advance task instructions, dACC may proactively modulate VTC until goal information becomes available. This performance-related difference in dACC *→* VTC SFC was no longer observed during the stim 2 window in late instruction trials (Extended Data Fig. 8), during which task instructions were available. This finding is consistent with dACC-VTC coupling being selectively engaged under instructional uncertainty.

We next asked whether strong dACC-VTC coupling was also observed under conditions of high cognitive demand. To test this, we compared SFC between pro and anti trials during the stim 1 window, noting that participants performed significantly worse and slower on anti trials than on pro trials (Fig. 1c), thereby indicating that anti trials were more cognitively demanding. SFC was significantly elevated in the theta-band (4-7 Hz) for anti trials compared to pro trials (Fig. 5e). Furthermore, when trials were separated by accuracy, theta-band coherence was significantly stronger for correct compared to incorrect responses in anti trials, whereas no such difference was observed for pro trials (Fig. 5f). These effects were confirmed using paired Wilcoxon signed-rank tests performed separately within each frequency band, with false discovery rate (FDR) correction applied across four frequency bands (theta 4-7 Hz; alpha 8-12 Hz; beta 12-32 Hz; gamma 32-80 Hz). After FDR correction, accuracy-related differences were significant only in the theta band for anti trials (*P* = 0.0368), with no significant effects observed in the alpha, beta, or gamma bands (all *P >* 0.7), nor in any frequency band for pro trials (all *P >* 0.45).

These findings suggest that dACC-VTC coupling increases not only during context uncertainty but also during high cognitive demand, and that stronger coupling supports successful performance in these conditions, specifically within the brief stim 1 window. Such rapid modulation of VTC by dACC may reflect a flexible top-down mechanism through which medial frontal cortices transiently synchronize sensory representations to optimize goal-directed performance.

## Discussion

We investigated how the timing of task instructions dynamically shapes sensory representations in the human brain using simultaneous single-neuron recordings across multiple cortical and medial temporal regions. We show that identical visual stimuli evoke distinct patterns of activity in VTC depending on whether task instructions preceded the onset of the stimuli or not, demonstrating that sensory representations in human VTC are flexibly and rapidly reorganized by instruction timing even when sensory input is held constant. To our knowledge, the current study provides the first human single-neuron and population-level evidence that the availability of task goals can dynamically reshape representations in a high-level visual area traditionally viewed as primarily sensory on a moment-by-moment timescale. At the same time, medial frontal regions, particularly dACC, carried robust signals related to instruction timing and exhibited frequency-specific, performance-predictive interactions with VTC, revealing a rapid, online top-down mechanism through which sensory population codes were modulated to support goal-directed computation. Importantly, this is the first evidence in humans that top-down interactions between medial frontal cortex and a high-level visual area are selectively strengthened under instructional uncertainty and increased cognitive demand in a manner that predicts behavioral performance.

Converging evidence from studies in animals suggests that top-down feedback from frontal and cingulate cortices modulates sensory processing through frequency-specific oscillatory coupling. Beta-band inter-areal synchronization has been linked to the maintenance of task sets and the transmission of predictive signals from higher-order areas to sensory cortices [27, 29, 30], whereas theta-band synchronization supports adaptive updating and cognitive control under uncertainty [31, 32]. The increase in dACC-VTC beta coherence we observed during the stimulus 1 window for late-instruction trials likely reflects top-down stabilization of sensory codes when goal information must be retroactively integrated with already-encoded stimuli. Notably, although explicit task instructions had not yet been provided at the onset of stimulus 1 in this condition, their absence itself appears to engage top-down control from dACC to structure sensory representations under uncertainty. Critically, stronger beta-band coupling during the stimulus 1 window predicted successful performance specifically when uncertainty was highest (in late instruction trials), suggesting that this mechanism is behaviorally consequential. These findings are aligned with previously described roles of beta-band coupling in mediating cognitive control [15, 33–35]. Conversely, elevated theta-band coupling during anti trials may index dynamic control processes required to apply inverted task rules (which are cognitively more demanding than pro trials). Similarly, theta enhancement also predicted correct performance specifically under high cognitive demand, supporting its role in coordinating flexible rule application. These frequency-dependent interactions are consistent with hierarchical predictive coding frameworks in which beta oscillations convey top-down predictions while theta oscillations coordinate cognitive control and error monitoring [36–39]. The dissociation between beta-mediated modulation during uncertainty and theta-mediated modulation during cognitive demand suggests that dACC employs frequency-multiplexed channels to flexibly coordinate with VTC. This architecture may enable the same circuit to simultaneously maintain stable sensory representations while allowing for rapid, context-dependent reconfiguration when task demands shift.

At the microcircuit level, feedback from medial frontal regions could engage specific classes of inhibitory interneurons in VTC to implement the context-dependent modulation we observed. In line with this possibility, prior studies in mice have demonstrated that top-down projections from higher-order cortical areas specifically engage vasoactive intestinal peptide (VIP)-expressing inhibitory neurons in visual cortex, which in turn inhibit somatostatin (SST)-expressing interneurons, resulting in disinhibition of pyramidal neurons [20, 21, 28]. These studies further show that such disinhibitory circuits are critical for enabling context-dependent modulation of visual responses, including task-related enhancement of behaviorally relevant features and suppression of irrelevant information. In our data, the sharpening of category representations in VTC when instructions preceded stimuli (Fig. 3) might reflect precisely this type of disinhibitory mechanism: dACC feedback may selectively release task-relevant VTC neurons from local inhibition, amplifying their responses while leaving the underlying categorical tuning intact. In addition, feedback could recruit parvalbumin (PV)-expressing interneurons to sharpen sensory representations through lateral inhibition, as has been observed during attention-demanding tasks in rodents [40, 41]. Testing these microcircuit hypotheses in humans remains challenging, but future studies combining laminar recordings with computational modeling could help disambiguate the specific inhibitory motifs underlying the top-down modulation we observed.

The robust dACC-VTC coupling we observed raises important questions about the anatomical pathways supporting this long-range communication. In non-human primates, anatomical tracing studies have revealed that anterior cingulate cortex sends direct feedback projections to inferotemporal cortex, terminating predominantly in superficial layers where they can modulate the activity of both excitatory and inhibitory neurons [42, 43]. These feedback projections follow a hierarchical gradient, with higher-order frontal regions preferentially targeting superficial layers of lower-order sensory areas, a pattern consistent with predictive coding models in which top-down predictions modulate sensory processing [44]. In humans, diffusion tensor imaging and tract-tracing studies have similarly identified white matter pathways connecting medial frontal cortex to ventral temporal regions, including connections through the inferior fronto-occipital fasciculus and the cingulum bundle [45, 46]. However, direct monosynaptic connections between dACC and VTC in humans may be sparse or nonexistent, raising the possibility that dACC modulation could also operate through polysynaptic routes involving intermediate relay stations such as the thalamus, posterior cingulate cortex, or lateral prefrontal regions [47–49]. The pulvinar nucleus of the thalamus represents a particularly intriguing candidate, as it receives inputs from prefrontal cortex and projects extensively to temporal visual areas, providing a potential substrate for rapid routing of contextual signals to sensory cortex [38, 47]. The pulvinar’s role in coordinating cortical communication through cortico-thalamo-cortical loops could enable dACC to dynamically gate sensory processing in VTC depending on behavioral demands [50, 51]. The specific contribution of monosynaptic versus polysynaptic pathways to the observed spike-field coupling remains an important question for future investigation, potentially addressable through combined recordings with electrical microstimulation or optogenetic perturbations in animal models.

Our findings have important implications for understanding cognitive flexibility and its breakdown in clinical populations. The dynamic reconfiguration of VTC representations we observed suggests that even canonical sensory areas are readily shaped by top-down goal states rather than operating as passive feature detectors. Importantly, the strength of dACC-VTC coupling predicted performance under conditions of high uncertainty and cognitive demand, suggesting that individual differences in this mechanism may underlie variability in cognitive control abilities. Dysfunction in such top-down modulation could contribute to cognitive inflexibility observed in neuropsychiatric disorders including schizophrenia, where patients show impaired context-dependent modulation of sensory processing [52], and autism spectrum disorder, where atypical sensory responses may reflect reduced top-down predictive signaling [53, 54]. Furthermore, neurodegenerative conditions such as Alzheimer’s disease, which preferentially affect frontal and temporal cortical regions, as well as age-related decline in prefrontal function, could weaken dACC-VTC coupling and contribute to reduced cognitive flexibility in older adults. Our findings thus provide a mechanistic framework for understanding how goal-directed modulation of sensory cortex supports adaptive behavior, and how disruptions to this mechanism may contribute to cognitive and perceptual deficits across clinical populations. While the present study focused specifically on how the timing of task instructions dynamically reorganizes sensory coding in VTC, the question of where and how instruction content itself is represented within this distributed network is addressed in a companion study using the same task framework.

## Acknowledgments

We are grateful to the staff of the epilepsy monitoring unit at Cedars-Sinai Medical Center, including the neuromonitoring staff, nursing staff, and epileptologists (Drs. Lisa Bateman and Jeffrey Chung), for their dedicated patient care and clinical support. We thank members of the Rutishauser laboratory, including Jonathan Daume, Hristos Courellis, Varun Wadia, Natasha Kurilenko, Jake Gavenas, and Daniel Deng, for their helpful discussions and feedback on the manuscript. We give special thanks to Varun Wadia and Hristos Courellis for their invaluable assistance with data collection for several patients. We also thank Jorge Aldana for assistance with computing resources. Above all, we are deeply grateful to the patients and their families for their willingness to participate in this research. This work was funded by the American Academy of Neurology (AAN) Resident Research Scholarship (to R.K.), the Simons Foundation’s SCGB program (to U.R.), and the BRAIN initiative through U01NS117839 (to U.R.). The funders had no role in study design, data collection and analysis, decision to publish, or preparation of the manuscript.

## Author contributions

N.R. and T.G.A. conceived the study. N.R., U.R. and T.G.A. designed the behavioral task. R.K. and T.G.A. collected the data. R.K. and T.G.A. performed data processing. R.K. performed formal analysis. S.C. localized electrodes and prepared the data release. R.K., N.R., and U.R. wrote the initial draft of the manuscript. T.G.A., C.M.R. and A.N.M. reviewed and edited the manuscript. N.R. and U.R. supervised the study. C.M.R. provided patient care and facilitated experiments. A.N.M. performed all surgical procedures. U.R. acquired funding.

## Methods

### Participants

Eleven adult patients (6 female; age range: 25 - 55 years; see Extended Data Table 1) with drug-resistant epilepsy participated in this study. All patients were undergoing stereotactic depth electrode implantation for seizure localization and potential surgical treatment at Cedars-Sinai Medical Center. Electrode placement was determined solely on clinical grounds by the clinical care team. Each patient provided written informed consent [55] to participate in research protocols approved by the Institutional Review Board of Cedars-Sinai Medical Center.

### Task design

Participants performed a flexible visual decision-making task designed to dissociate sensory encoding from task-related contextual modulation. The task was implemented in MATLAB using Psychtoolbox (v3.0; [56]). Each participant completed one recording session consisting of four blocks of 48 trials each. The blocks alternated between early and late instruction conditions.

In the early instruction condition, each trial began with a 1-s fixation period followed by a 2.5-s instruction period. Two color images, drawn from one of four visual categories (cars, animals, fruits, and human faces), were then presented sequentially for 1 s each. Based on the instruction, participants indicated whether the first or second image satisfied the rule by pressing a corresponding button. The mapping between the first/second choice and left/right response buttons was randomized across trials to prevent motor preparation biases. In the late instruction condition, each trial also began with a 1-s fixation period but was followed by two sequential images before the 2.5-s instruction period, after which participants made their choice.

There were two types of instructions: pro and anti. In pro trials, participants followed the rule as stated (e.g., “Choose the most colorful car”), whereas in anti trials they applied the opposite rule (e.g., “Do not choose the most colorful car”). Because of their double-negative structure, anti trials imposed greater cognitive demands than pro trials.

All conditions were balanced across blocks such that each image appeared twice, as the first and second stimulus, and was presented equally often under early and late instruction conditions.

### Electrophysiology and spike sorting

Intracranial recordings were obtained at Cedars-Sinai Medical Center from bilateral hybrid depth electrodes containing eight microwires each (AdTech Medical) [57, 58]. Electrode implantation sites were determined exclusively by clinical needs for seizure localization and included the ventral temporal cortex (VTC), dorsal anterior cingulate cortex (dACC), pre-supplementary motor area (preSMA), ventromedial prefrontal cortex (vmPFC), hippocampus (HIPP), and amygdala (AMY). See Extended Data Table 1. Broadband extracellular signals (0.1–9,000 Hz) were continuously recorded from each microwire at a sampling rate of 32 kHz using the ATLAS system (Neuralynx Inc.). Recordings were locally referenced within each electrode bundle to a designated low-impedance microwire.

Electrode localization was performed using pre-operative T1-weighted magnetic resonance imaging (MRI) scans and post-operative computed tomography (CT) scans. Image processing and alignment were conducted using FreeSurfer (v5.3.0 and v7.4.1) and Advanced Normalization Tools (ANTs v2.1) [59]. Post-operative CT scans were rigidly co-registered to the corresponding pre-operative MRI scans. Each MRI scan was mapped to the MNI152-registered CIT168 probabilistic atlas using the symmetric normalization (SyN) algorithm implemented in ANTs. The resulting affine and nonlinear transformations were then applied to the co-registered CT scan. Finally, electrode localization was performed in MNI space using 3D Slicer (v5.0.3) and Freeview (FreeSurfer). Electrode contacts were visually inspected and confirmed to be within gray matter.

Spike detection and sorting were performed offline using the semi-automated template-matching algorithm OSort (v4.1 [60]). The raw broadband signal was band-pass filtered between 300 and 3,000 Hz, and candidate spikes were detected using an adaptive threshold. Putative single units were isolated based on waveform shape, refractory period violations, and cluster separation. Units that did not meet these isolation criteria were excluded. Only patients with at least one well-isolated single neuron in one of the regions of interest were included in the analyses.

### LFP preprocessing

All local field potential (LFP) analyses were performed on broadband signals (0.1 - 9,000 Hz, sampled at 32 kHz) recorded from the microwires of the hybrid depth electrodes. To prevent contamination of low-frequency activity by action-potential (spiking) artifacts, we removed spike waveforms by linearly interpolating the raw signal from −1 to +2 ms around each detected spike time on that microwire. This interpolation was applied across the continuous broadband recording prior to downsampling. Because the same spike can occasionally appear on multiple nearby microwires, interpolation was performed across all wires within the same bundle whenever a spike was detected on any one of them.

After spike removal, signals underwent band-stop filtering to remove 60 Hz line noise and its harmonics up to 180 Hz using a zero-phase Butterworth filter (2 Hz bandwidth around each harmonic). The data were then low-pass filtered at 500 Hz (8th-order Butterworth, zero-phase) and downsampled to 1 kHz using MATLAB (MathWorks, R2023a). All filtering and downsampling steps were implemented using zero-phase (forward-reverse) filtering to avoid phase distortions in the LFP signal.

Following these preprocessing steps, each trial and channel were individually reviewed by visual inspection. Trials containing interictal epileptiform discharges, large-amplitude movement artifacts, amplifier saturation, or abnormally low signal variance were excluded from further analysis. Channels showing persistent interictal discharges or excessive noise throughout the session were removed. On average, 60.02 *±* 69.27 trials (31.3% of the data) were excluded per microwire during this artifact rejection process. Only artifact-free trials and channels were included in subsequent analyses.

### Single-neuron selectivity analysis

Neuronal selectivity for stimulus category, instruction timing, and their interaction was quantified using a two-way analysis of variance (ANOVA) performed independently for each neuron. Analyses were conducted separately for responses to the first and second stimuli (stim 1 and stim 2). For VTC neurons, firing rates were calculated within a 0.1 - 1.0 s window following stimulus onset (t=0 is stimulus onset). For non-VTC regions (dorsal anterior cingulate cortex, pre-supplementary motor area, ventromedial prefrontal cortex, hippocampus, and amygdala), the analysis window was shifted to 0.2 - 1.0 s after stimulus onset to account for the longer onset latencies in higher-order areas [61].

For each neuron, trial-by-trial firing rates were entered into a two-way 2 *×* 4 ANOVA with instruction timing (early vs. late) and stimulus category (cars, animals, fruits, faces) as fixed factors, including the interaction term. Neurons were classified as category-selective or instruction-selective based on significant main effects (*P <* 0.05, uncorrected). Neurons showing a significant main effect of category were labeled category-selective, and those with a significant main effect of instruction timing were labeled instruction-selective. Neurons showing significant main effects for both factors were grouped as jointly selective.

The proportion of neurons in each class was computed for each brain region. For population-level summaries (Fig. 2d and Fig. 4a), neurons were counted once per region, and chance-level proportions (5%) were estimated via shuffling of category and instruction labels across trials.

### Population decoding analysis

Population decoding was performed using a pseudo-population approach, in which single-trial firing rates from all isolated neurons recorded within a given brain region were aggregated across participants, as has been done in prior studies [62–64]. This approach allowed us to estimate population-level discriminability despite the limited number of simultaneously recorded neurons per participant. For each neuron, firing rates were averaged within the stimulus analysis window (0.1-1.0 s after stimulus onset for VTC; 0.2-1.0 s for non-VTC regions).

Decoding analyses were implemented in MATLAB (MathWorks, R2023a) using a linear support vector machine (SVM) classifier (fitcecoc function with default parameters). For category decoding, data were balanced across the four stimulus categories (cars, animals, fruits, faces), with four trials per category used for training and four for testing (16 trials total per split). This process was repeated for 100 random iterations, and decoding accuracy was averaged across iterations.

For instruction-order decoding, analyses were conducted separately within each stimulus category to ensure that sensory information was held constant. Within each category, four trials from each instruction condition (early vs. late) were used for training and four for testing (eight trials total per split). Trial assignments were randomized across 100 iterations, and the mean decoding accuracy across iterations was used as the final measure.

All decoding analyses were performed on datasets in which the number of trials was explicitly matched across conditions to avoid bias from unequal trial counts. Decoding performance was evaluated on held-out test data in each iteration, and reported accuracies represent mean cross-validated decoding performance across iterations for each brain region.

For comparing SVM decoding accuracy across brain areas (Fig. 4b,d), the number of neurons used for decoding was fixed to the number of isolated neurons from the region with the smallest sample size (dACC, 74 neurons). For regions containing more than 74 neurons, 74 neurons were randomly subsampled at each iteration to ensure matched population sizes across regions.

### State-space analysis using CEBRA

To visualize the low-dimensional structure of population activity, we used Consistent Embeddings of high-dimensional Recordings using Auxiliary variables (CEBRA; [25]). Unlike linear dimensionality reduction methods such as principal component analysis (PCA), CEBRA learns nonlinear latent embeddings that preserve the temporal continuity and intrinsic geometry of neural population dynamics.

We used an unsupervised version of CEBRA, fitting the model exclusively to neural activity without auxiliary behavioral or task labels. The model was trained on concatenated single-trial firing rates from all neurons within each region, using the same stimulus-aligned analysis windows defined above. Prior to training, we performed a grid search to identify optimal model parameters (output dimensions, time offsets, hidden units, and temperature) that minimized the InfoNCE contrastive loss. The final model was trained using the “offset10-model” architecture with a batch size of 1024, learning rate of 3 *×*10*^−^*^4^, temperature of 1.1, and cosine distance metric. The model was conditioned on time with a time offset of 5, and trained for up to 3000 iterations with 64 hidden units and an output dimensionality of 6. All training was performed using the official CEBRA Python package (v0.3.0) with GPU acceleration enabled when available.

Each trained model produced low-dimensional embeddings for every trial, which were used to visualize neural trajectories for early and late instruction conditions. To quantify the separation between trajectories, we computed the Euclidean distance between the mean latent trajectories of the two conditions at each time point. These distances provided a measure of representational divergence in the latent space, with shuffled-label controls used to confirm that observed trajectory differences were above chance. To ensure that our results were not dependent on a single model initialization, we trained 10 independent CEBRA models using identical hyperparameters but different random seeds. For each brain region, low-dimensional trajectories and Euclidean distance time courses of early and late instruction conditions were computed separately within each trained model and then averaged across models to obtain the final trajectory estimates and distance curves.

### Spike-field coherence

To quantify the strength of phase coupling between neuronal spiking activity and LFPs, we computed spike–field coherence (SFC) [65] during the stim 1 window (0.1–1.0 s after stimulus onset for VTC; 0.2–1.0 s for non-VTC regions). Analyses were restricted to neuron–LFP channel pairs recorded within the same hemisphere.

Instantaneous phase information from LFPs was extracted using complex Morlet wavelets spanning 2 - 150 Hz in 40 logarithmically spaced frequency steps. The number of cycles for each wavelet increased logarithmically from 3 to 10 across the frequency range to ensure adequate spectral and temporal resolution. For each neuron-LFP pair, we computed the phase of the LFP at each spike time and quantified SFC as the mean vector length (MVL) of the spike-phase distribution across all spikes and trials.

To control for differences in trial counts and spike rate across conditions, the number of trials and the spike-time distribution were matched between the two conditions being compared (e.g., early vs. late instruction, correct vs. incorrect, pro vs. anti). For each matched dataset, SFC was computed using the procedure described above. This process was repeated 100 iterations, and the mean SFC across iterations was used as the condition-specific estimate.

To assess statistical significance, we generated a null distribution of SFC values by shuffling spike times within each trial while preserving the overall firing rate profile. For each iteration, spike times during the stim 1 window were shuffled, and SFC was recomputed using the same procedure. This shuffling process was repeated 100 times, and the resulting surrogate distribution was used to compute a z-scored SFC for the oringinal data.

### Statistics

Throughout the study, statistical analyses were performed using non-parametric methods unless otherwise stated. Pairwise comparisons between independent samples used the Wilcoxon rank-sum test, and paired comparisons used the Wilcoxon signed-rank test. All tests were two-sided unless noted, and *P <* 0.05 was considered significant. When applicable, trial counts and spike numbers were matched across conditions to avoid biases arising from unequal sample sizes. Additional task- or analysis-specific statistical procedures (e.g., ANOVA for single-neuron selectivity, Kruskal–Wallis tests with Dunn’s correction for comparing decoding accuracy across regions, and cluster-based bootstrap testing for time-resolved effects) are described below and in the corresponding sections.

#### Cluster-based bootstrap testing

To identify time periods showing significant differences between two conditions, we used a nonparametric cluster-based bootstrap procedure. For each comparison, we first computed the observed time-pointwise difference between conditions. We then generated a null distribution by randomly permuting condition labels across trials (or across neurons for pseudo-population analyses) and recomputing the time-pointwise difference on each iteration (1,000 bootstrap iterations). Significant time points were identified by thresholding the observed difference against the 95th percentile of the null distribution (one-sided). Contiguous suprathreshold time points were grouped into clusters, and a cluster-level statistic was computed as the sum of the absolute differences within each cluster. The same procedure was applied to each bootstrap iteration to obtain a null distribution of cluster-level statistics. Clusters in the observed data whose cluster statistic exceeded the 95th percentile of the bootstrap null distribution were considered significant (*P <* 0.05, cluster-level corrected). All significant clusters are indicated in figures by thick horizontal black bars.

## Code Availability

The code for the analyses performed in this work will be made publicly available upon acceptance.

## Data Availability

Data used in this study will be made publicly available upon acceptance.

## Extended Data Tables

**Extended Data Table 1.**
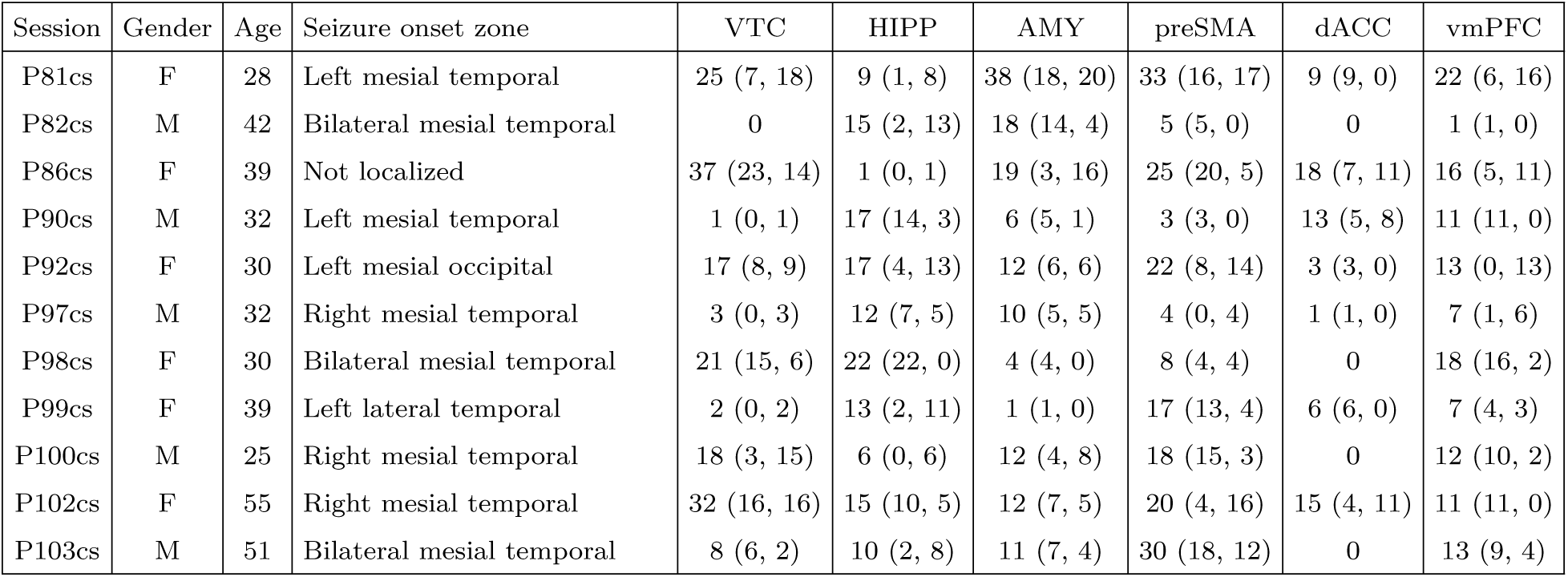
Summary of patient demographics and single-neuron recordings across brain regions. Each entry indicates the total number of well-isolated neurons obtained from spike sorting. Values in parentheses correspond to left and right hemisphere counts, respectively.

| Session | Gender | Age | Seizure onset zone | VTC | HIPP | AMY | preSMA | dACC | vmPFC |
| --- | --- | --- | --- | --- | --- | --- | --- | --- | --- |
| P81cs | F | 28 | Left mesial temporal | 25 (7, 18) | 9 (1, 8) | 38 (18, 20) | 33 (16, 17) | 9 (9, 0) | 22 (6, 16) |
| P82cs | M | 42 | Bilateral mesial temporal | 0 | 15 (2, 13) | 18 (14, 4) | 5 (5, 0) | 0 | 1 (1, 0) |
| P86cs | F | 39 | Not localized | 37 (23, 14) | 1 (0, 1) | 19 (3, 16) | 25 (20, 5) | 18 (7, 11) | 16 (5, 11) |
| P90cs | M | 32 | Left mesial temporal | 1 (0, 1) | 17 (14, 3) | 6 (5, 1) | 3 (3, 0) | 13 (5, 8) | 11 (11, 0) |
| P92cs | F | 30 | Left mesial occipital | 17 (8, 9) | 17 (4, 13) | 12 (6, 6) | 22 (8, 14) | 3 (3, 0) | 13 (0, 13) |
| P97cs | M | 32 | Right mesial temporal | 3 (0, 3) | 12 (7, 5) | 10 (5, 5) | 4 (0, 4) | 1 (1, 0) | 7 (1, 6) |
| P98cs | F | 30 | Bilateral mesial temporal | 21 (15, 6) | 22 (22, 0) | 4 (4, 0) | 8 (4, 4) | 0 | 18 (16, 2) |
| P99cs | F | 39 | Left lateral temporal | 2 (0, 2) | 13 (2, 11) | 1 (1, 0) | 17 (13, 4) | 6 (6, 0) | 7 (4, 3) |
| P100cs | M | 25 | Right mesial temporal | 18 (3, 15) | 6 (0, 6) | 12 (4, 8) | 18 (15, 3) | 0 | 12 (10, 2) |
| P102cs | F | 55 | Right mesial temporal | 32 (16, 16) | 15 (10, 5) | 12 (7, 5) | 20 (4, 16) | 15 (4, 11) | 11 (11, 0) |
| P103cs | M | 51 | Bilateral mesial temporal | 8 (6, 2) | 10 (2, 8) | 11 (7, 4) | 30 (18, 12) | 0 | 13 (9, 4) |

**Extended Data Table 2.** Example task instructions for Pro and Anti conditions.

| **Pro condition** | **Anti condition** |
| --- | --- |
| Choose the older car | Do not select the older car |
| Select the image with fewer objects | Do not select the image with fewer objects |
| Choose the more colorful animal | Avoid the more colorful animal |
| Select the less modern vehicle | Do not select the less modern vehicle |
| Choose the image with more animals | Do not select the image with more animals |
| Select the vehicle with more colors | Do not select the vehicle with more colors |
| Choose the younger face | Avoid the younger face |
| Select the creature with fewer colors | Do not select the creature with fewer colors |
| Choose the image with more creatures | Do not select the image with more creatures |
| Select the more modern vehicle | Do not select the more modern vehicle |
| Choose the image with fewer items | Do not select the image with fewer items |
| Select the more colorful car | Avoid the more colorful car |

**Extended Data Table 3.** Pairwise post-hoc comparisons of instruction-order decoding accuracy across regions (related to Extended Data Fig. 4). Differences across the six regions were assessed using a Kruskal-Wallis nonparametric ANOVA, followed by Dunn’s multiple comparisons test. The table reports all pairwise contrasts for both the stim 1 and stim 2 windows, including group indices, mean rank differences, confidence intervals, and adjusted *P*-values.

| Stim 1 |  |  |  |  |  |
| --- | --- | --- | --- | --- | --- |
| Region 1 | Region 2 | Mean diff | CI low | CI high | *P*-value |
| VTC | HIPP | -183.61 | -112.28 | -40.94 | 0.0001 |
| VTC | AMY | -34.14 | 37.20 | 108.53 | 0.8693 |
| VTC | preSMA | -228.67 | -157.33 | -85.99 | < 0.0001 |
| VTC | dACC | -353.24 | -281.91 | -210.57 | < 0.0001 |
| VTC | vmPFC | -142.86 | -71.53 | -0.19 | 0.0488 |
| HIPP | AMY | 78.13 | 149.48 | 220.89 | < 0.0001 |
| HIPP | preSMA | -116.39 | -45.06 | 26.28 | 0.6310 |
| HIPP | dACC | -240.97 | -169.63 | -98.29 | < 0.0001 |
| HIPP | vmPFC | -29.58 | 40.75 | 112.09 | 0.7724 |
| AMY | preSMA | -194.37 | -124.55 | -63.28 | < 0.0001 |
| AMY | dACC | -265.86 | -194.56 | -123.17 | < 0.0001 |
| AMY | vmPFC | -80.95 | -8.72 | 37.28 | 0.0001 |
| preSMA | dACC | -189.46 | -120.18 | -37.38 | 0.0001 |
| preSMA | vmPFC | 14.47 | 85.85 | 157.14 | 0.0064 |
| dACC | vmPFC | 139.04 | 210.38 | 281.72 | < 0.0001 |
| Stim 2 |  |  |  |  |  |
| Region 1 | Region 2 | Mean diff | CI low | CI high | *P*-value |
| VTC | HIPP | -353.90 | -282.41 | -210.91 | < 0.0001 |
| VTC | AMY | -448.01 | -376.56 | -305.03 | < 0.0001 |
| VTC | preSMA | -198.15 | -126.66 | -55.16 | < 0.0001 |
| VTC | dACC | -425.43 | -353.94 | -282.45 | < 0.0001 |
| VTC | vmPFC | -222.03 | -150.54 | -79.04 | < 0.0001 |
| HIPP | AMY | 165.61 | 94.12 | 22.62 | 0.0017 |
| HIPP | preSMA | -84.26 | 155.75 | 227.24 | < 0.0001 |
| HIPP | dACC | -143.03 | -71.54 | -0.04 | 0.0497 |
| HIPP | vmPFC | 160.37 | 31.87 | 103.36 | < 0.0001 |
| AMY | preSMA | 178.37 | 249.87 | 321.36 | 0.9936 |
| AMY | dACC | -48.91 | 22.76 | 94.07 | < 0.0001 |
| AMY | vmPFC | 154.49 | 225.98 | 297.47 | < 0.0001 |
| preSMA | dACC | -298.78 | -227.29 | -155.79 | < 0.0001 |
| preSMA | vmPFC | -95.38 | -23.89 | 47.61 | 0.9974 |
| dACC | vmPFC | 131.91 | 203.40 | 274.89 | < 0.0001 |

**Extended Data Table 4.** Pairwise Dunn-test comparisons across early/late conditions in all regions during stim 1 window (related to Extended Data Fig. 5a). Each row lists a pairwise contrast between regional conditions (e.g., VTC Early vs. HIPP Late), including the confidence-interval lower bound, mean rank difference, confidence-interval upper bound, and Dunn corrected *P*-value.

| Region 1 | Region 2 | CI low | Mean diff | CI high | *P*-value |
| --- | --- | --- | --- | --- | --- |
| VTC Early | VTC Late | -41.7522 | 121.3750 | 284.5022 | 0.5610 |
| VTC Early | HIPP Early | 243.6328 | 406.7600 | 569.8872 | < 0.0001 |
| VTC Early | HIPP Late | 145.7528 | 308.8800 | 472.0072 | < 0.0001 |
| VTC Early | AMY Early | 216.2428 | 379.3700 | 542.4972 | < 0.0001 |
| VTC Early | AMY Late | 444.2728 | 607.4000 | 770.5272 | < 0.0001 |
| VTC Early | preSMA Early | 342.4028 | 505.5300 | 668.6572 | < 0.0001 |
| VTC Early | preSMA Late | 542.4128 | 705.5400 | 868.6672 | < 0.0001 |
| VTC Early | dACC Early | 527.2078 | 690.3350 | 853.4622 | < 0.0001 |
| VTC Early | dACC Late | 472.2078 | 635.3350 | 798.4622 | < 0.0001 |
| VTC Early | vmPFC Early | 447.9878 | 611.1150 | 774.2422 | < 0.0001 |
| VTC Early | vmPFC Late | 226.8328 | 389.9600 | 553.0872 | < 0.0001 |
| VTC Late | HIPP Early | 122.2578 | 285.3850 | 448.5122 | < 0.0001 |
| VTC Late | HIPP Late | 24.3778 | 187.5050 | 350.6322 | 0.0074 |
| VTC Late | AMY Early | 94.8678 | 257.9950 | 421.1222 | < 0.0001 |
| VTC Late | AMY Late | 322.8978 | 486.0250 | 649.1522 | < 0.0001 |
| VTC Late | preSMA Early | 221.0278 | 384.1550 | 547.2822 | < 0.0001 |
| VTC Late | preSMA Late | 421.0378 | 584.1650 | 747.2922 | < 0.0001 |
| VTC Late | dACC Early | 405.8328 | 568.9600 | 732.0872 | < 0.0001 |
| VTC Late | dACC Late | 350.8328 | 513.9600 | 677.0872 | < 0.0001 |
| VTC Late | vmPFC Early | 326.6128 | 489.7400 | 652.8672 | < 0.0001 |
| VTC Late | vmPFC Late | 105.4578 | 268.5850 | 431.7122 | < 0.0001 |
| HIPP Early | HIPP Late | -261.0072 | -97.8800 | 65.2472 | 0.9477 |
| HIPP Early | AMY Early | -190.5172 | -27.3900 | 135.7372 | 1.0000 |
| HIPP Early | AMY Late | 37.5128 | 200.6400 | 363.7672 | 0.0024 |
| HIPP Early | preSMA Early | -64.3572 | 98.7700 | 261.8972 | 0.9405 |
| HIPP Early | preSMA Late | 135.6528 | 298.7800 | 461.9072 | < 0.0001 |
| HIPP Early | dACC Early | 120.4478 | 283.5750 | 446.7022 | < 0.0001 |
| HIPP Early | dACC Late | 65.4478 | 228.5750 | 391.7022 | 0.0002 |
| HIPP Early | vmPFC Early | 41.2278 | 204.3550 | 367.4822 | 0.0017 |
| HIPP Early | vmPFC Late | -179.9272 | -16.8000 | 146.3272 | 1.0000 |
| HIPP Late | AMY Early | -92.6372 | 70.4900 | 233.6172 | 1.0000 |
| HIPP Late | AMY Late | 135.3928 | 298.5200 | 461.6472 | < 0.0001 |
| HIPP Late | preSMA Early | 33.5228 | 196.6500 | 359.7772 | 0.0034 |
| HIPP Late | preSMA Late | 233.5328 | 396.6600 | 559.7872 | < 0.0001 |
| HIPP Late | dACC Early | 218.3278 | 381.4550 | 544.5822 | < 0.0001 |
| HIPP Late | dACC Late | 163.3278 | 326.4550 | 489.5822 | < 0.0001 |
| HIPP Late | vmPFC Early | 139.1078 | 302.2350 | 465.3622 | < 0.0001 |
| HIPP Late | vmPFC Late | -82.0472 | 81.0800 | 244.2072 | 0.9986 |
| AMY Early | AMY Late | 64.9028 | 228.0300 | 391.1572 | 0.0002 |
| AMY Early | preSMA Early | -36.9672 | 126.1600 | 289.2872 | 0.4618 |

**Extended Data Table 4 | Pairwise Dunn-test comparisons across early/late conditions in all regions during stim 1 window (related to Extended Data Fig.**
| Region 1 | Region 2 | CI low | Mean diff | CI high | *P*-value |
| --- | --- | --- | --- | --- | --- |
| AMY Early | preSMA Late | 163.0428 | 326.1700 | 489.2972 | < 0.0001 |
| AMY Early | dACC Early | 147.8378 | 310.9650 | 474.0922 | < 0.0001 |
| AMY Early | dACC Late | 92.8378 | 255.9650 | 419.0922 | < 0.0001 |
| AMY Early | vmPFC Early | 68.6178 | 231.7450 | 394.8722 | 0.0001 |
| AMY Early | vmPFC Late | -152.5372 | 10.5900 | 173.7172 | 1.0000 |
| AMY Late | preSMA Early | -264.9972 | -101.8700 | 61.2572 | 0.9100 |
| AMY Late | preSMA Late | -64.9872 | 98.1400 | 261.2672 | 0.9457 |
| AMY Late | dACC Early | -80.1922 | 82.9350 | 246.0622 | 0.9976 |
| AMY Late | dACC Late | -135.1922 | 27.9350 | 191.0622 | 1.0000 |
| AMY Late | vmPFC Early | -159.4122 | 3.7150 | 166.8422 | 1.0000 |
| AMY Late | vmPFC Late | -380.5672 | -217.4400 | -54.3128 | 0.0005 |
| preSMA Early | preSMA Late | 36.8828 | 200.0100 | 363.1372 | 0.0025 |
| preSMA Early | dACC Early | 21.6778 | 184.8050 | 347.9322 | 0.0092 |
| preSMA Early | dACC Late | -33.3222 | 129.8050 | 292.9322 | 0.3910 |
| preSMA Early | vmPFC Early | -57.5422 | 105.5850 | 268.7122 | 0.8624 |
| preSMA Early | vmPFC Late | -278.6972 | -115.5700 | 47.5572 | 0.6831 |
| preSMA Late | dACC Early | -178.3322 | -15.2050 | 147.9222 | 1.0000 |
| preSMA Late | dACC Late | -233.3322 | -70.2050 | 92.9222 | 1.0000 |
| preSMA Late | vmPFC Early | -257.5522 | -94.4250 | 68.7022 | 0.9700 |
| preSMA Late | vmPFC Late | -478.7072 | -315.5800 | -152.4528 | < 0.0001 |
| dACC Early | dACC Late | -218.1272 | -55.0000 | 108.1272 | 1.0000 |
| dACC Early | vmPFC Early | -242.3472 | -79.2200 | 83.9072 | 0.9992 |
| dACC Early | vmPFC Late | -463.5022 | -300.3750 | -137.2478 | < 0.0001 |
| dACC Late | vmPFC Early | -187.3472 | -24.2200 | 138.9072 | 1.0000 |
| dACC Late | vmPFC Late | -408.5022 | -245.3750 | -82.2478 | < 0.0001 |
| vmPFC Early | vmPFC Late | -384.2822 | -221.1550 | -58.0278 | 0.0003 |

**Extended Data Table 5.** Pairwise Dunn-test comparisons across early/late conditions in all regions during stim 2 window (related to Extended Data Fig. 5b). Each row lists a pairwise contrast between regional conditions (e.g., VTC Early vs. HIPP Late), including the confidence-interval lower bound, mean rank difference, confidence-interval upper bound, and Dunn corrected *P*-value.

| Region 1 | Region 2 | CI low | Mean diff | CI high | *P*-value |
| --- | --- | --- | --- | --- | --- |
| VTC Early | VTC Late | -263.8578 | -100.6350 | 62.5878 | 0.9238 |
| VTC Early | HIPP Early | 282.3722 | 445.5950 | 608.8178 | < 0.0001 |
| VTC Early | HIPP Late | 275.4422 | 438.6650 | 601.8878 | < 0.0001 |
| VTC Early | AMY Early | 280.2522 | 443.4750 | 606.6978 | < 0.0001 |
| VTC Early | AMY Late | -20.4678 | 142.7550 | 305.9778 | 0.1954 |
| VTC Early | preSMA Early | 159.9972 | 323.2200 | 486.4428 | < 0.0001 |
| VTC Early | preSMA Late | 446.9772 | 610.2000 | 773.4228 | < 0.0001 |
| VTC Early | dACC Early | 399.0022 | 562.2250 | 725.4478 | < 0.0001 |
| VTC Early | dACC Late | 182.1872 | 345.4100 | 508.6328 | < 0.0001 |
| VTC Early | vmPFC Early | 368.1972 | 531.4200 | 694.6428 | < 0.0001 |
| VTC Early | vmPFC Late | 214.2872 | 377.5100 | 540.7328 | < 0.0001 |
| VTC Late | HIPP Early | 383.0072 | 546.2300 | 709.4528 | < 0.0001 |
| VTC Late | HIPP Late | 376.0772 | 539.3000 | 702.5228 | < 0.0001 |
| VTC Late | AMY Early | 380.8872 | 544.1100 | 707.3328 | < 0.0001 |
| VTC Late | AMY Late | 80.1672 | 243.3900 | 406.6128 | < 0.0001 |
| VTC Late | preSMA Early | 260.6322 | 423.8550 | 587.0778 | < 0.0001 |
| VTC Late | preSMA Late | 547.6122 | 710.8350 | 874.0578 | < 0.0001 |
| VTC Late | dACC Early | 499.6372 | 662.8600 | 826.0828 | < 0.0001 |
| VTC Late | dACC Late | 282.8222 | 446.0450 | 609.2678 | < 0.0001 |
| VTC Late | vmPFC Early | 468.8322 | 632.0550 | 795.2778 | < 0.0001 |
| VTC Late | vmPFC Late | 314.9222 | 478.1450 | 641.3678 | < 0.0001 |
| HIPP Early | HIPP Late | -170.1528 | -6.9300 | 156.2928 | 1.0000 |
| HIPP Early | AMY Early | -165.3428 | -2.1200 | 161.1028 | 1.0000 |
| HIPP Early | AMY Late | -466.0628 | -302.8400 | -139.6172 | < 0.0001 |
| HIPP Early | preSMA Early | -285.5978 | -122.3750 | 40.8478 | 0.5414 |
| HIPP Early | preSMA Late | 1.3822 | 164.6050 | 327.8278 | 0.0452 |
| HIPP Early | dACC Early | -46.5928 | 116.6300 | 279.8528 | 0.6626 |
| HIPP Early | dACC Late | -263.4078 | -100.1850 | 63.0378 | 0.9282 |
| HIPP Early | vmPFC Early | -77.3978 | 85.8250 | 249.0478 | 0.9950 |
| HIPP Early | vmPFC Late | -231.3078 | -68.0850 | 95.1378 | 1.0000 |
| HIPP Late | AMY Early | -158.4128 | 4.8100 | 168.0328 | 1.0000 |
| HIPP Late | AMY Late | -459.1328 | -295.9100 | -132.6872 | < 0.0001 |
| HIPP Late | preSMA Early | -278.6678 | -115.4450 | 47.7778 | 0.6871 |
| HIPP Late | preSMA Late | 8.3122 | 171.5350 | 334.7578 | 0.0269 |
| HIPP Late | dACC Early | -39.6628 | 123.5600 | 286.7828 | 0.5166 |
| HIPP Late | dACC Late | -256.4778 | -93.2550 | 69.9678 | 0.9758 |
| HIPP Late | vmPFC Early | -70.4678 | 92.7550 | 255.9778 | 0.9779 |
| HIPP Late | vmPFC Late | -224.3778 | -61.1550 | 102.0678 | 1.0000 |
| AMY Early | AMY Late | -463.9428 | -300.7200 | -137.4972 | < 0.0001 |
| AMY Early | preSMA Early | -283.4778 | -120.2550 | 42.9678 | 0.5862 |

**Extended Data Table 5 | Pairwise Dunn-test comparisons across early/late conditions in all regions during stim 2 window (related to Extended Data Fig.**
| Region 1 | Region 2 | CI low | Mean diff | CI high | *P*-value |
| --- | --- | --- | --- | --- | --- |
| AMY Early | preSMA Late | 3.5022 | 166.7250 | 329.9478 | 0.0386 |
| AMY Early | dACC Early | -44.4728 | 118.7500 | 281.9728 | 0.6181 |
| AMY Early | dACC Late | -261.2878 | -98.0650 | 65.1578 | 0.9467 |
| AMY Early | vmPFC Early | -75.2778 | 87.9450 | 251.1678 | 0.9918 |
| AMY Early | vmPFC Late | -229.1878 | -65.9650 | 97.2578 | 1.0000 |
| AMY Late | preSMA Early | 17.2422 | 180.4650 | 343.6878 | 0.0133 |
| AMY Late | preSMA Late | 304.2222 | 467.4450 | 630.6678 | < 0.0001 |
| AMY Late | dACC Early | 256.2472 | 419.4700 | 582.6928 | < 0.0001 |
| AMY Late | dACC Late | 39.4322 | 202.6550 | 365.8778 | 0.0020 |
| AMY Late | vmPFC Early | 225.4422 | 388.6650 | 551.8878 | < 0.0001 |
| AMY Late | vmPFC Late | 71.5322 | 234.7550 | 397.9778 | 0.0001 |
| preSMA Early | preSMA Late | 123.7572 | 286.9800 | 450.2028 | < 0.0001 |
| preSMA Early | dACC Early | 75.7822 | 239.0050 | 402.2278 | 0.0001 |
| preSMA Early | dACC Late | -141.0328 | 22.1900 | 185.4128 | 1.0000 |
| preSMA Early | vmPFC Early | 44.9772 | 208.2000 | 371.4228 | 0.0012 |
| preSMA Early | vmPFC Late | -108.9328 | 54.2900 | 217.5128 | 1.0000 |
| preSMA Late | dACC Early | -211.1978 | -47.9750 | 115.2478 | 1.0000 |
| preSMA Late | dACC Late | -428.0128 | -264.7900 | -101.5672 | < 0.0001 |
| preSMA Late | vmPFC Early | -242.0028 | -78.7800 | 84.4428 | 0.9993 |
| preSMA Late | vmPFC Late | -395.9128 | -232.6900 | -69.4672 | 0.0001 |
| dACC Early | dACC Late | -380.0378 | -216.8150 | -53.5922 | 0.0005 |
| dACC Early | vmPFC Early | -194.0278 | -30.8050 | 132.4178 | 1.0000 |
| dACC Early | vmPFC Late | -347.9378 | -184.7150 | -21.4922 | 0.0094 |
| dACC Late | vmPFC Early | 22.7872 | 186.0100 | 349.2328 | 0.0084 |
| dACC Late | vmPFC Late | -131.1228 | 32.1000 | 195.3228 | 1.0000 |
| vmPFC Early | vmPFC Late | -317.1328 | -153.9100 | 9.3128 | 0.0961 |

## Extended Data Figures

**Extended Data Fig. 1.**
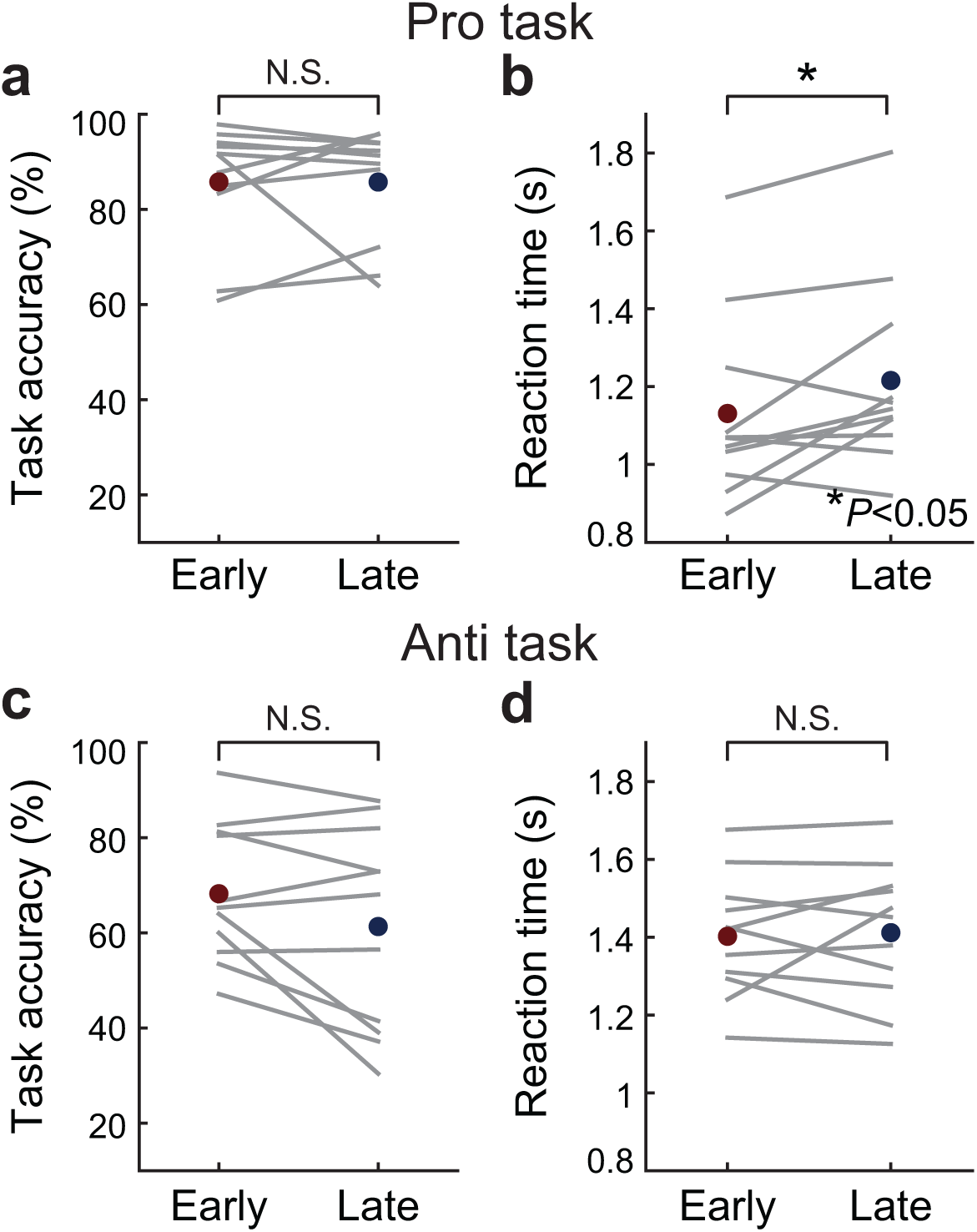
Behavioral performance for early versus late instruction trials within pro and anti conditions. **a**, Pro task. Task accuracy (left) and reaction time (right) shown for early and late instruction trials. Each gray line represents an individual participant, and colored points denote group means (blue: early; red: late). Accuracy did not differ between early and late pro trials. However, reaction times were significantly faster for early compared to late pro trials (*P <* 0.05, one-sided Wilcoxon signed-rank test). **b**, Anti task. Task accuracy (left) and reaction time (right) for early versus late instruction trials. Neither accuracy nor reaction time differed significantly between early and late anti trials.

**Extended Data Fig. 2.**
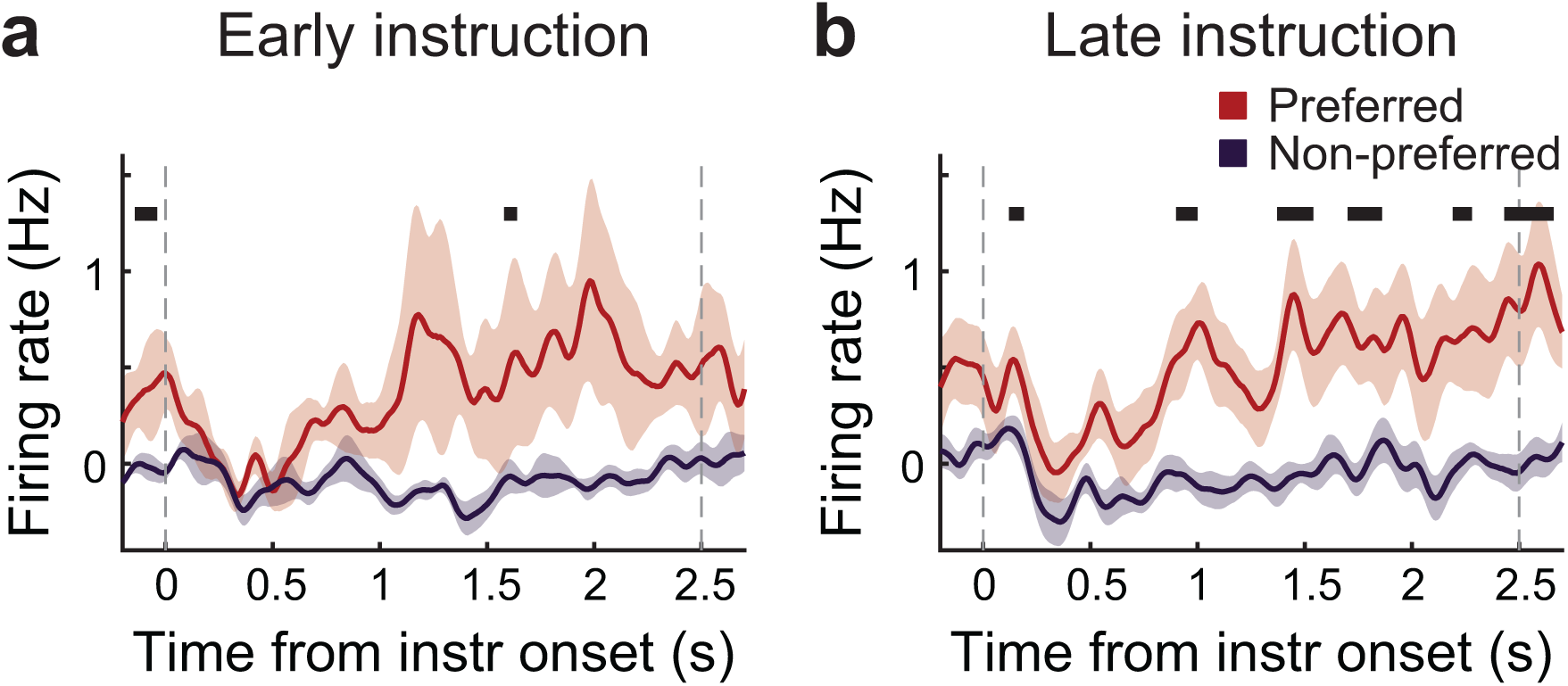
Subset of category-selective VTC neurons show instruction-dependent firing-rate modulation. **a**, average firing rates (mean *±* s.e.m.) across the 7 VTC category-selective neurons (from 3 participants) that were identified, using a bootstrap null distribution of preferred-versus-non-preferred firing-rate differences, as exhibiting significantly elevated activity in the instruction window during early-instruction trials. **b**, average firing rates across the 13 VTC category-selective neurons (from 5 participants) that met the same bootstrap-null criterion for the late-instruction condition. Between the two instruction-timing conditions, 5 neurons contributed to both groups. Thick horizontal black bars indicate time periods of significant preferred-versus-non-preferred differences based on the cluster-based bootstrap procedure (*P <* 0.05, cluster-level corrected). Vertical dashed lines mark instruction onset and offset, respectively.

**Extended Data Fig. 3.**
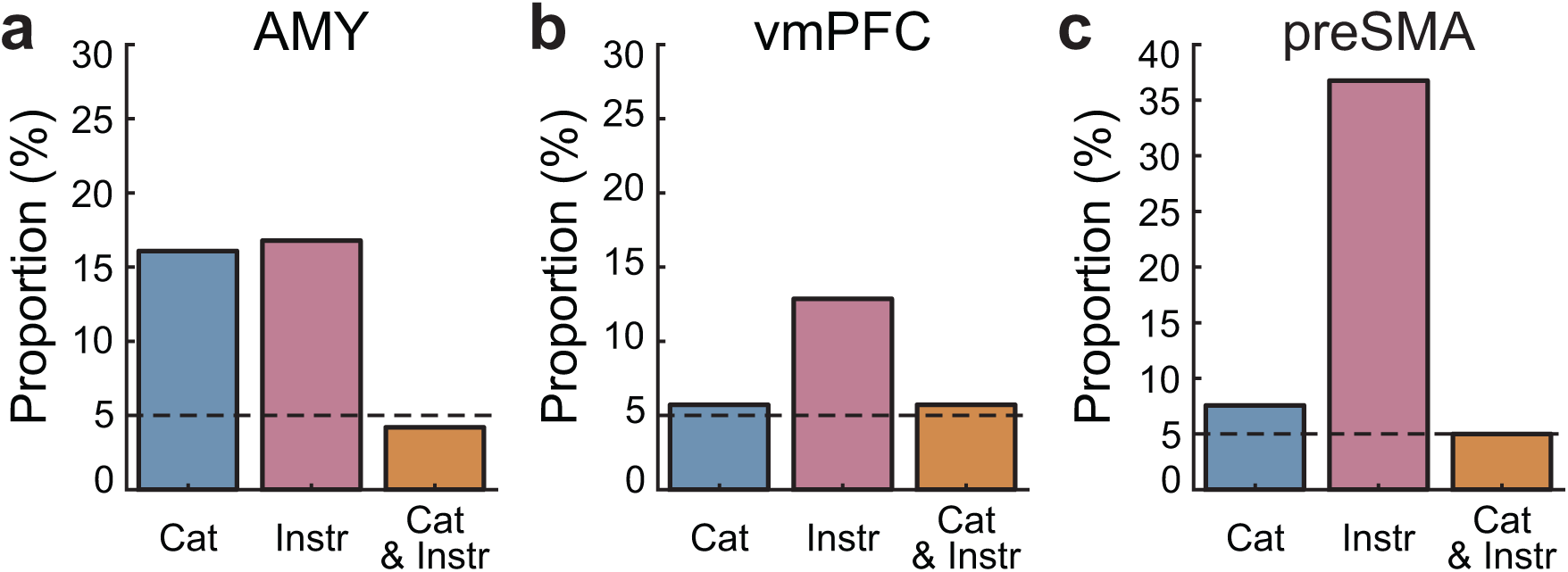
Category-, instruction-, and jointly selective neurons in additional higher-order regions. Proportion of neurons exhibiting selectivity for stimulus category (Cat; magenta), instruction timing (Instr; teal), or both factors (Cat+Instr; orange) based on a two-way ANOVA (category *×* instruction timing) in (**a**) amygdala (AMY), (**b**) ventromedial prefrontal cortex (vmPFC), and (**c**) pre-supplementary motor area (preSMA). Dashed line indicates the 5% chance level.

**Extended Data Fig. 4.**
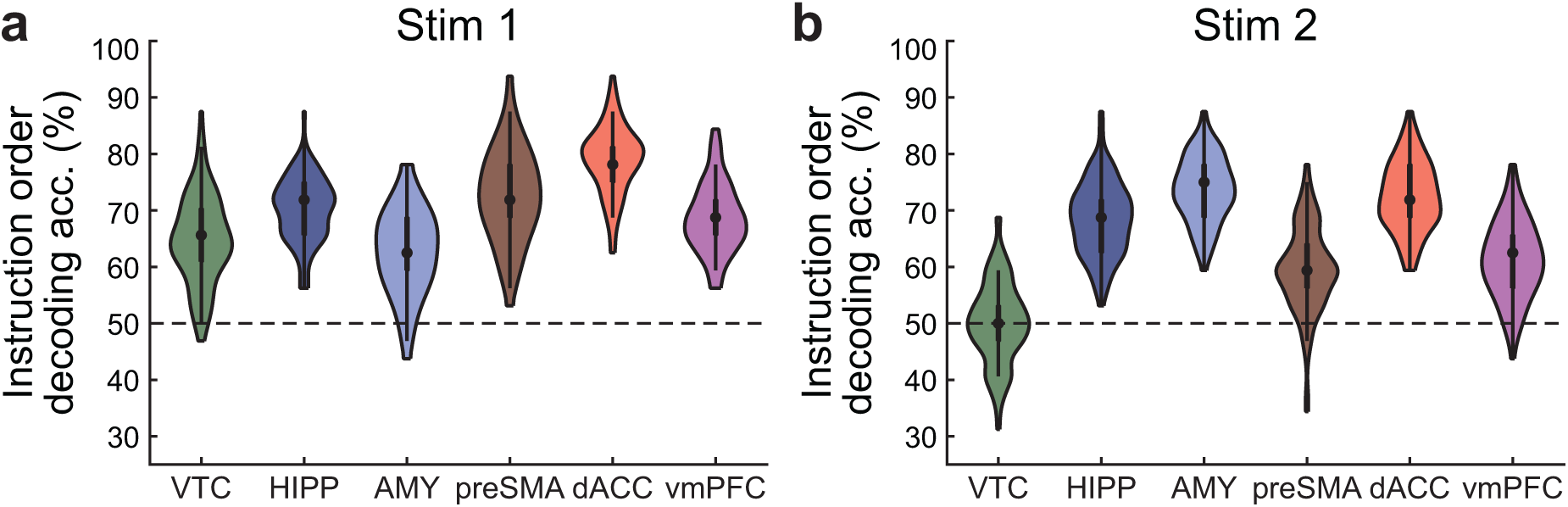
Instruction-order decoding accuracy across brain regions. Violin plots show linear SVM decoding accuracy for distinguishing early versus late instruction trials during the stim 1 (**a**) and stim 2 (**b**) windows across six regions: ventral temporal cortex (VTC), hippocampus (HIPP), amygdala (AMY), pre-supplementary motor area (preSMA), dorsal anterior cingulate cortex (dACC), and ventromedial prefrontal cortex (vmPFC). White dots denote median decoding accuracy, and the dashed horizontal line indicates chance level (50%). Kruskal–Wallis tests were performed across regions, followed by pairwise post-hoc Dunn’s multiple comparisons tests to identify significant differences (results reported in Extended Data Table 3.

**Extended Data Fig. 5.**
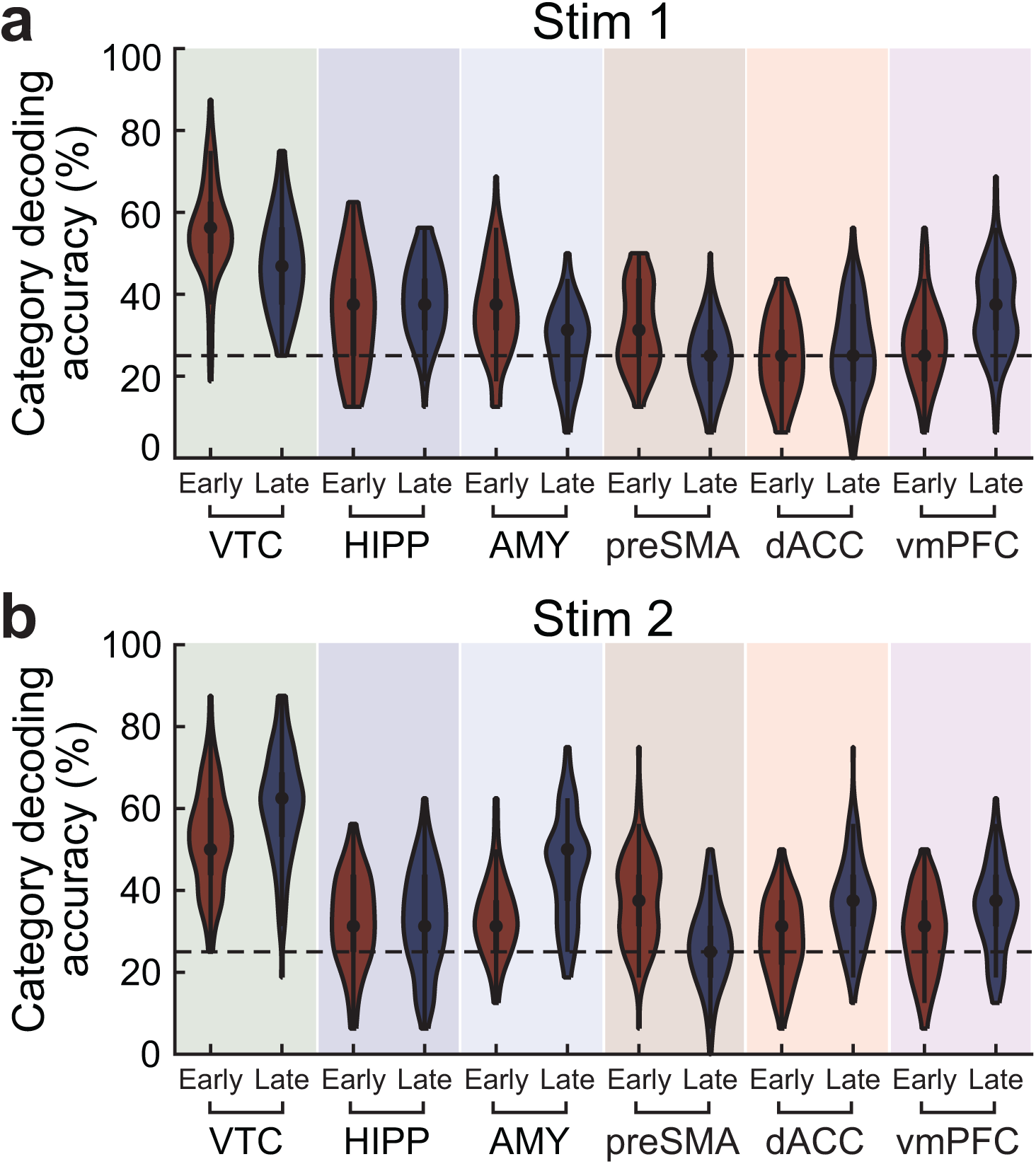
Category decoding accuracy across brain regions for stim 1 and stim 2 windows. Violin plots show linear SVM decoding accuracy for decoding stimulus category during the stim 1 (**a**) and stim 2 (**b**) windows across six regions: ventral temporal cortex (VTC), hippocampus (HIPP), amygdala (AMY), pre-supplementary motor area (preSMA), dorsal anterior cingulate cortex (dACC), and ventromedial prefrontal cortex (vmPFC). Early and late instruction trials are plotted separately for each region. White dots denote median decoding accuracy, and the dashed horizontal line marks chance level (25%). Kruskal–Wallis tests were performed across regions, followed by pairwise post-hoc Dunn’s multiple comparisons tests to identify significant differences (results reported in Extended Data Table 4–5).

**Extended Data Fig. 6.**
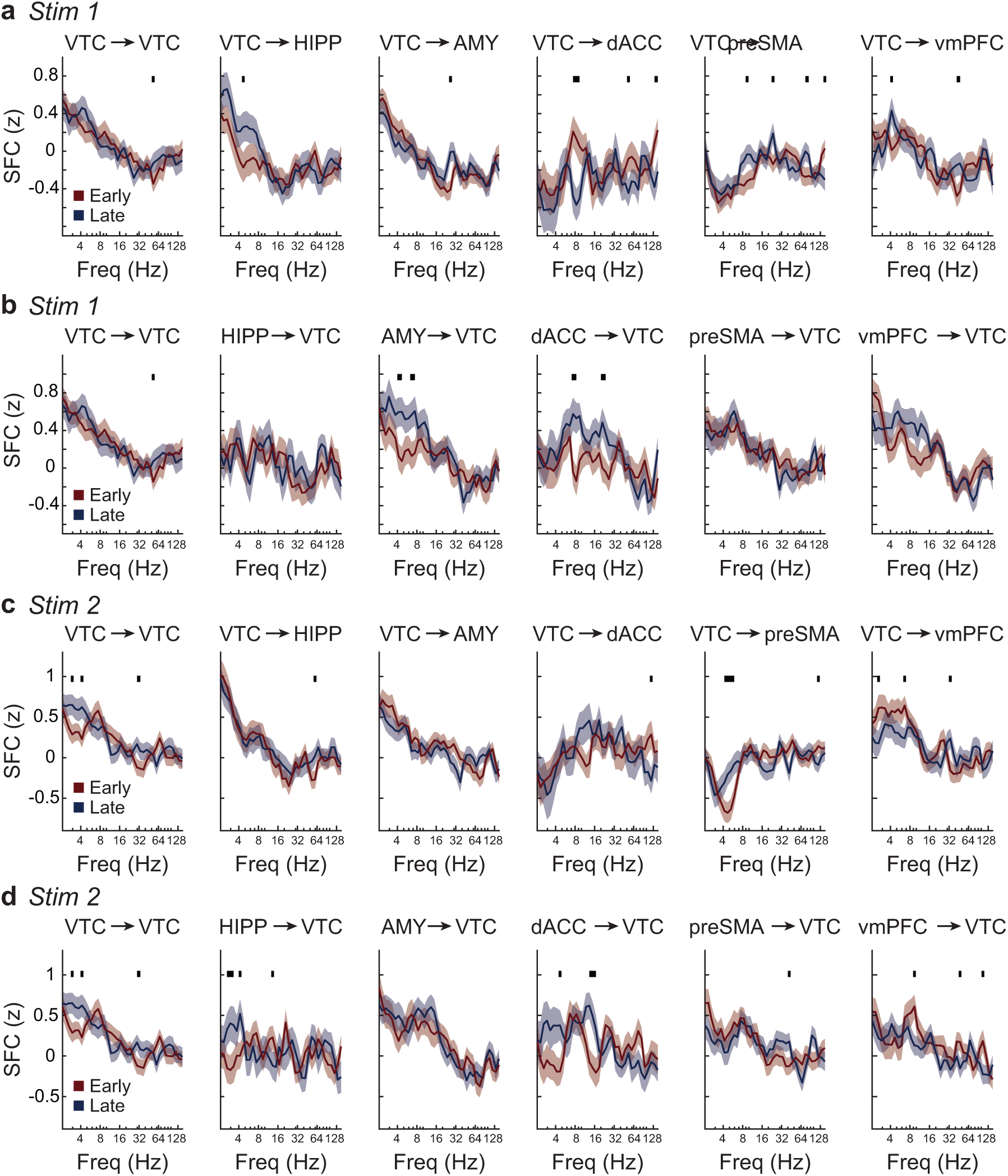
Spike-field coherence (SFC) differences between early and late instruction trials across VTC–network pairs. a-b, Stim 1 window. Population-averaged SFC (z-scored) is shown for early (dark blue) and late (dark red) instruction trials for all VTC-involving pairs, plotted separately for VTC as the LFP region (**a**) and VTC as the spike region (**b**). **c-d**, Stim 2 window. Same analyses as in **a** and **b**, computed during the stim 2 window. Shaded regions denote mean *±* s.e.m. across neurons. Thick horizontal black bars indicate time periods of significant preferred-versus-non-preferred differences based on the cluster-based bootstrap procedure (*P <* 0.05, cluster-level corrected).

**Extended Data Fig. 7.**
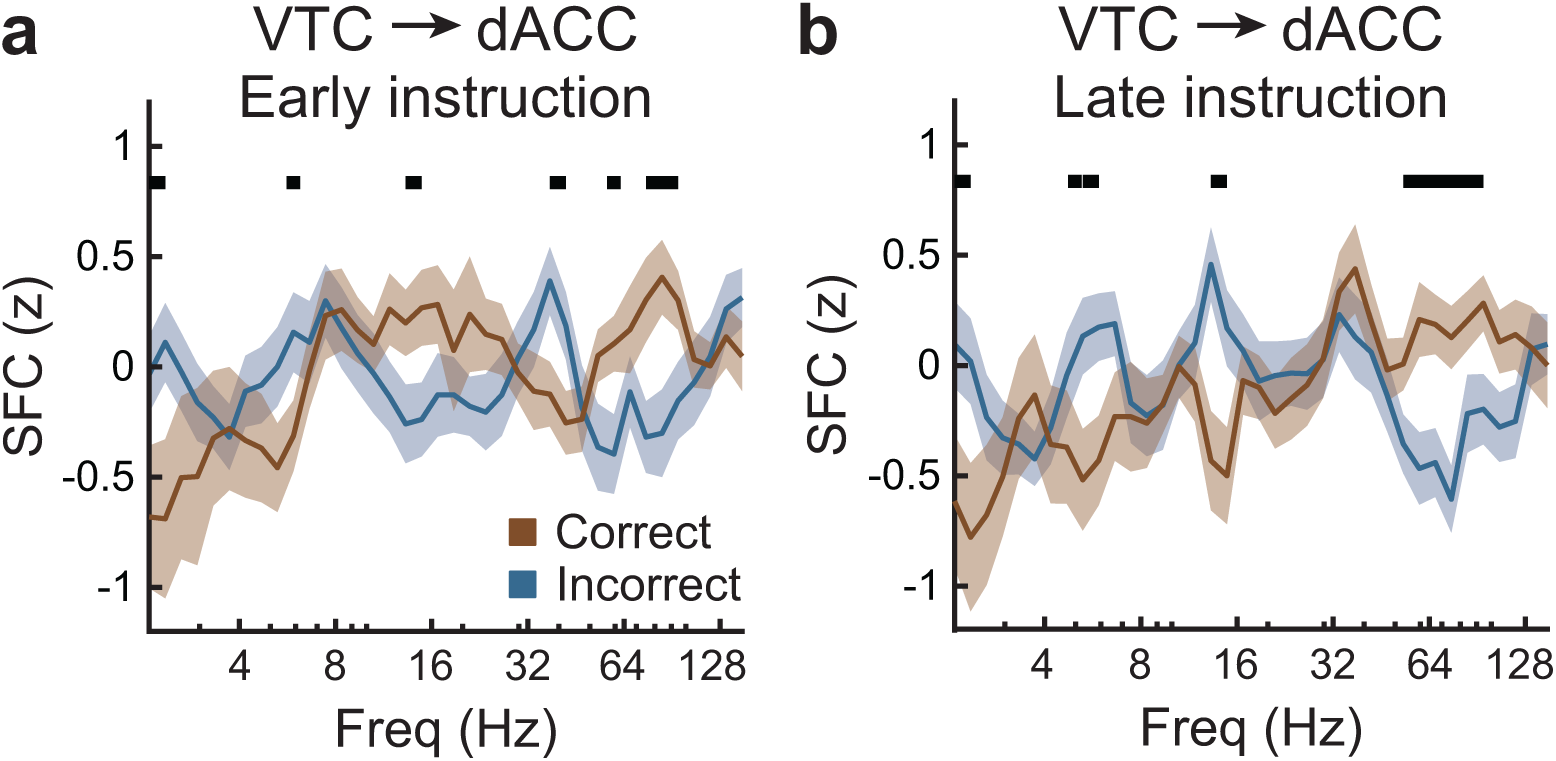
Spike-field coherence (SFC) differences between correct and incorrect trials for VTC LFP and dACC spikes. SFC (z-scored) is shown for dACC spikes phase-locked to VTC LFP signals during early (**a**) and late (**b**) instruction trials. Orange and blue traces denote correct and incorrect trials, respectively; shaded regions indicate mean *±* s.e.m. Thick horizontal black bars indicate time periods of significant preferred-versus-non-preferred differences based on the cluster-based bootstrap procedure (*P <* 0.05, cluster-level corrected).

**Extended Data Fig. 8.**
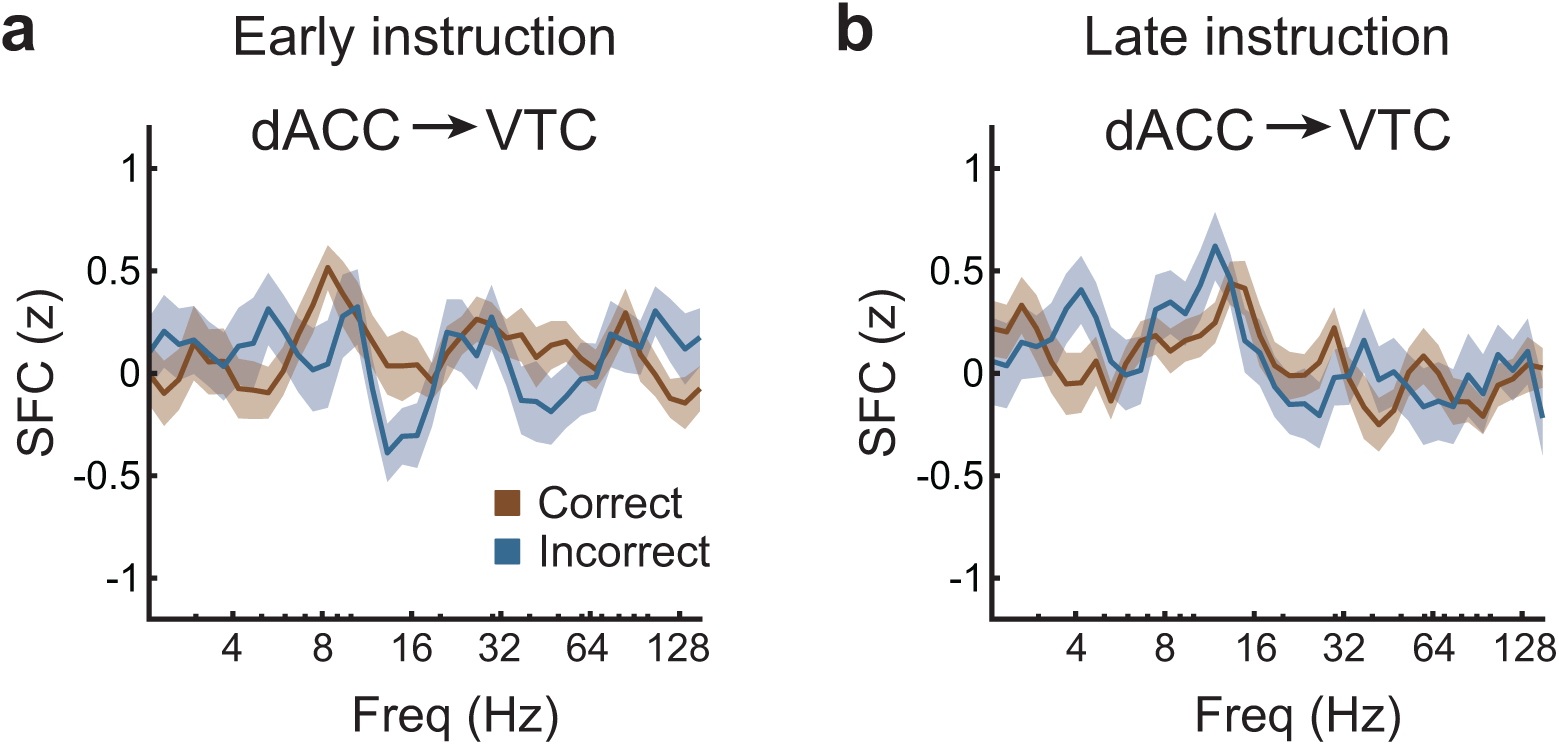
Spike-field coherence (SFC) differences between correct and incorrect trials for VTC spikes and dACC LFP during stim 2 windows. SFC (z-scored) is shown for VTC spikes phase-locked to dACC LFP signals during stim 2 window for early (**a**) and late (**b**) instruction conditions. Orange and blue traces denote correct and incorrect trials, respectively; shaded regions indicate mean *±* s.e.m.

